# RASGRP4 is a key factor in KRAS activation mediated by SOS in Y1 mouse tumor cell line

**DOI:** 10.1101/2025.07.23.663512

**Authors:** Fabio Montoni, Rosangela A. M. Wailemann, Thompson E. P. Torres, Katia A. M. Torres, Cecília S. Fonseca, Marcelo S. Reis, Hugo A. Armelin

## Abstract

The Y1 mouse adrenocortical carcinoma cell line presents amplification of the KRas oncogene and high-basal levels of KRAS-GTP mediated by the GEF SOS. In this research, we developed a dynamic model based on ordinary differential equations of the KRAS-GTP activation mediated by SOS in Y1 cells, which showed that SOS only is not sufficient to reach the high-basal levels of KRAS-GTP experimentally observed for this cell line. Interestingly, a modification in this system, which added another GEF in the model, made the model reach the expected levels of KRAS activation, leading to the hypothesis that there was a missing element in this system. To find this missing element, a PCR panel of RasGEFs was performed and the GEF Rasgrp4 was found highly expressed in parental Y1 cell lines, indicating that this was the missing element in the system. Finally, tumor growth assays in Balb/c-NUDE mice with the Y1 cell versus RASGRP4 CRISPR depleted Y1 cells, showed reduced tumor growth and frequency for the RASGRP4 depleted cells.

## Introduction

The Y1 cell line is a mouse derived adrenocortical carcinoma, an established model for the study of ACTH receptor-mediated steroidogenesis and signaling (1, 2). They present amplification of the KRas oncogene (3), with high basal-levels of KRAS-GTP, even in absence of external stimulation (4). In addition, Fibroblast Growth Factor 2 (FGF2), known for its role in neurogenesis, morphogenesis, wound healing (5), impairs the cell cycle in G2/M, restraining the proliferation of Y1 cell line via ERK pathway, leading to cell death (4, 6). These characteristics make this cell line relevant to study the MAPKs (RAS-RAF-MEK-ERK) pathway, which plays a key role in a number of cancers (7–9). The tumorigenicity observed in this cell line is a result of a deregulation of the switch mechanism that is controlled by the small GTPases (GEFs and GAPs), mediated by the GEF SOS (10, 11), resulting in the aforementioned high KRAS-GTP basal-levels. Small GTPases of the RAS superfamily (also called G-Proteins) are part of the GTP-binding proteins, which function as molecular switches, ranging from the GDP-bound inactive state and GTP-bound active state. This switching mechanism is tightly regulated by two types of accessory proteins: GTPase-activating proteins (GAPs), which promote the hydrolysis of GTP to GDP, switching RAS to its inactive state, and guanine nucleotide exchange factors (GEFs), which facilitate the release of GDP, allowing GTP to bind and activate RAS and consequently their downstream kinases (12, 13). It is also crucial to note that in the Y1 cell line, the amplification of the Kras gene itself is insufficient to drive tumorigenicity, as well as the KRAS activation mediated by SOS. The combination of these two factors must occur to consequently trigger the ERK signaling cascade that underlies this phenotype. Despite there is some level of understanding of the SOS mediation of this mechanism, much from the entangled interaction of the other GEFs is yet to be learned (14). To better understand this complex interaction, we have developed a dynamic model based on ordinary differential equations of the KRAS-GTP activation mediated by SOS in Y1 cells, which pointed to the need for an additional unknown factor. After careful investigation using classic molecular biology techniques and validation through mouse tumor assay, we report RASGRP4 as a novel necessary GEF to drive tumorigenicity in the Y1 cells, where its absence prevents the tumor growth in Balb/c-NUDE mice. These findings point to the need for greater attention to the GEFs as a whole, a rationale that can be applied in other clinical relevant cancer lineages.

## Results

### ODE modeling of the steady-state of KRAS activation in Y1 cells mediated by SOS displays unexpected instability

To construct the dynamic model of KRAS activation in Y1 cells, we chose to use the species and reactions previously described by Jayaji Das *et al*. in 2009 (7), which takes into account the allosteric regulation of SOS by RAS-GDP and RAS-GTP, that sets up a positive feedback (Figure 1). This model derived the following system of ODEs:

**Fig. 1.**
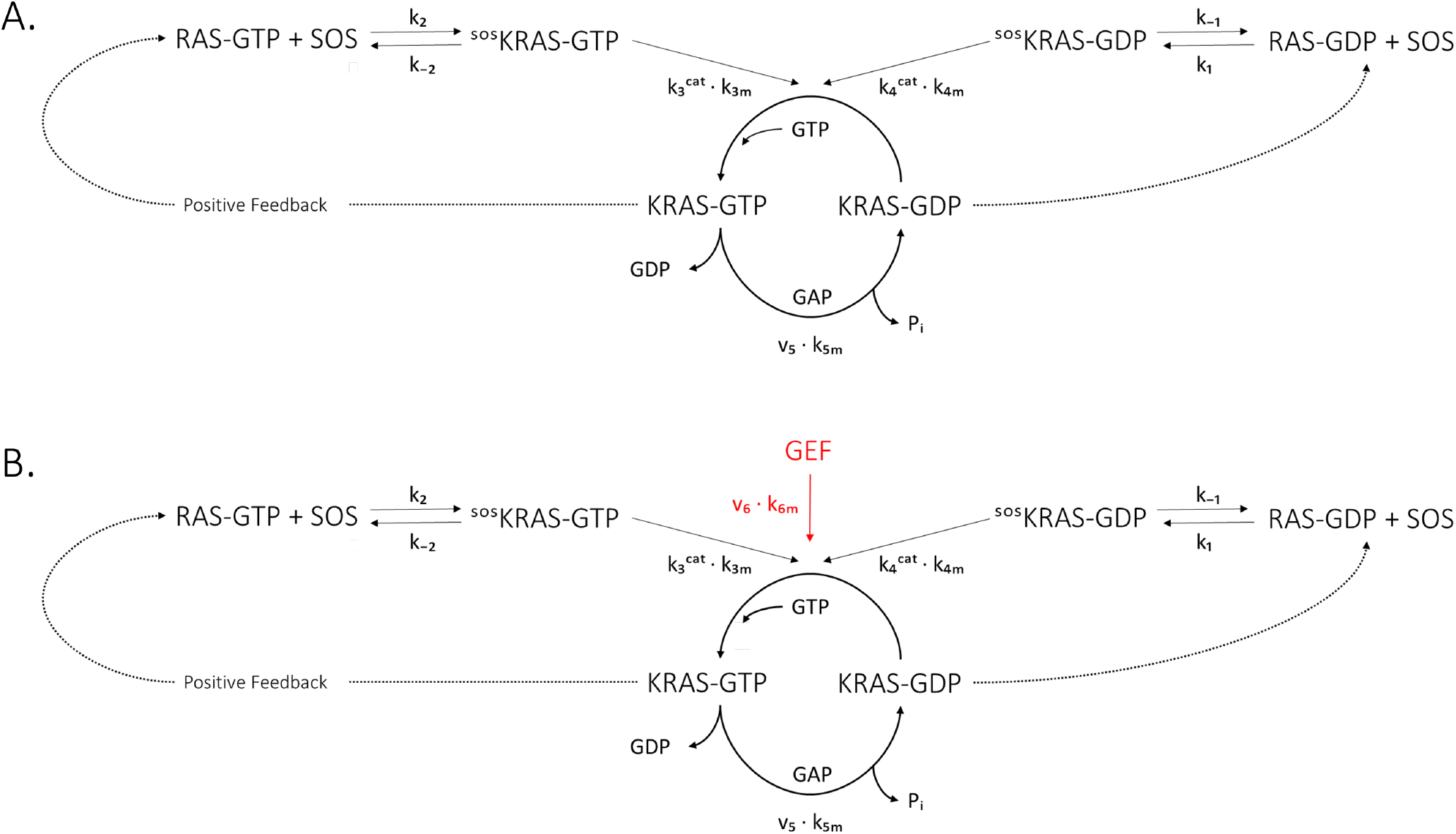
Species and chemical reactions that constitute the RAS molecular switch. Reaction rate constants are indicated next to them. Ras alternates between an active state (RAS-GTP), catalyzed by GEFs such as SOS, and an inactive state (RAS-GDP), catalyzed by GAPs. SOS is allosterically regulated by both RAS-GTP and RAS-GDP, which leads to an elevation of its catalytic activity. As allosteric regulation by RAS-GTP increases the catalytic activity of SOS by 75-fold (compared to a 15-fold increase provided by the same regulation by RAS-GDP), the molecular switch presents a positive feedback. **A**: RAS molecular switch with only SOS as GEF. **B**:The same switch with the inclusion of an additional GEF, which does not present positive feedback.

Where SOS-RAS-GDP and SOS-RAS-GTP denote SOS regulated allosterically by RAS-GDP and RAS-GTP, respectively. The rate constants were defined with experimentally measured values found in the literature (7).

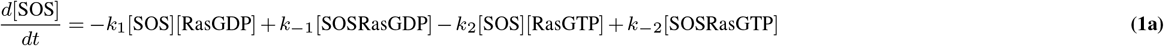

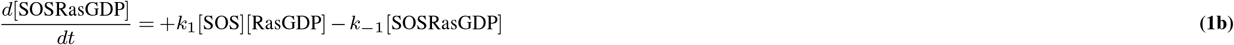

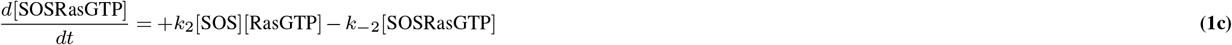

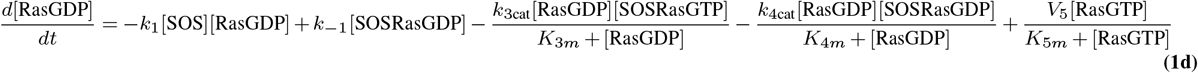

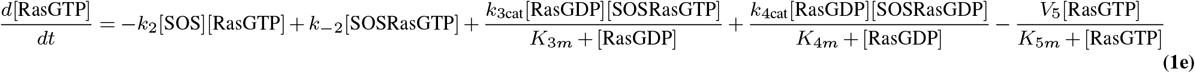

**Equation 1**: ODE modeling of the steady-state of KRAS activation in Y1 cells mediated by SOS. The Kinetic parameters of this model can be accessed in the Table 3.

Thus, we explored the newly defined model for different total concentrations of KRAS and SOS, increasing the former to up to 20 times the experimentally measured concentration in HeLa cells (7), in an attempt to mimic the high basal levels of KRAS-GTP observed in Y1 cells. However, we observed that for different total concentrations of KRAS, it was not possible to find a total SOS value that could explain the constitutively high basal levels of KRAS-GTP observed in Y1 cells. The system either entered a stationary state with negligible levels of KRAS-GTP or all KRAS remained in the KRAS-GTP form, thus yielding results inconsistent with those experimentally found in Y1 cells. Therefore, we speculated that the dynamic model was probably missing critical reactions to explain the functioning of the KRAS molecular switch in Y1 cells. Since our group reported years ago the involvement of GEFs in maintaining high basal levels of KRAS-GTP in Y1 cells (15), we decided to include an additional GEF in the dynamic model, whose presence in this cell line was still unknown; this was done by adding the term:

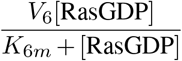

to the right side of equation (1e) of the ODE system, and the same term with a negative sign to the right side of equation (1d). With these modifications, it was possible to obtain a stable high basal level of KRAS-GTP. It is worth noting that the switch maintained the ability to alternate between “off” and “on”, although it lost the bistability observed in the first trial (Figure 2). The kinetic parameters can be seen on Table 3.

**Fig. 2.**
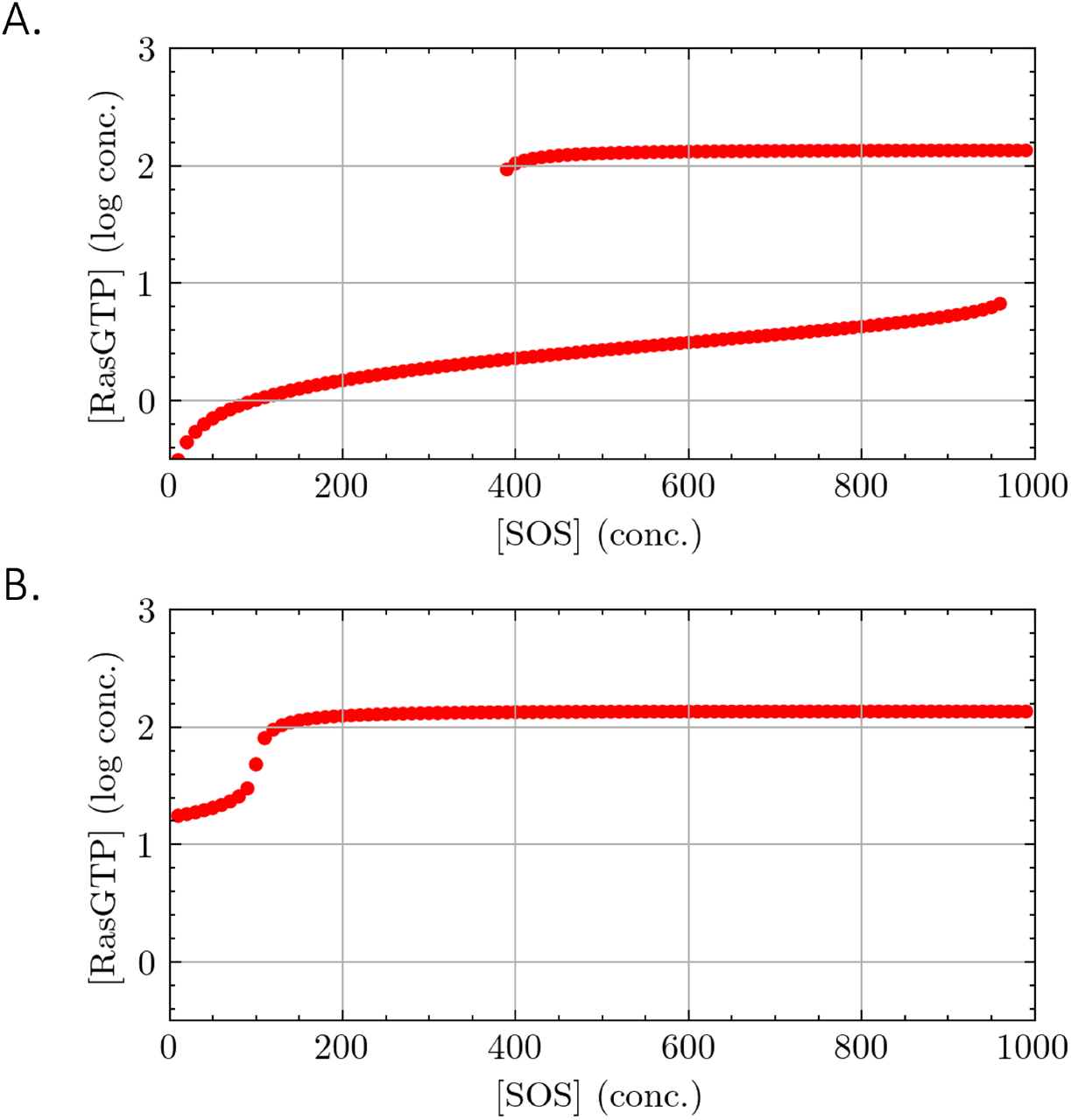
Elevated basal levels of RAS-GTP dependent on another GEF besides SOS. For a given total RAS concentration, states were computed for different SOS concentrations, taking the RAS-GTP concentration as the output reading. Data are presented on a log scale; units correspond to the original linear values. **A**: To calculate these steady states, the ODE system was used, counting only SOS as GEF **B**: ODE system including an additional GEF.

### PCR analysis of Y1 GEFs sheds the light for the unknown factor

In deprived Y1 cells, Figure 3A shows the expression levels of each GEF relative to the level of Sos1, where Rasgrp4 was the predominant GEF in parental cells, being 8 and 10 times more expressed than Grf2 and Sos1. Figure 3B shows the levels of each GEF in all lineages and growth conditions evaluated, always relative to the level of the respective Ras GEF in starved Y1 cells. The highlight in these gene expression results is that Rasgrp4, the predominant GEF in Y1 cells, is absent in all Y1-FD lines. Therefore, high basal levels of KRAS-GTP are likely found in parental Y1 cells depending concomitantly on high expression of total Kras and high levels of Rasgrp4 expression. This result prompted us to conclude that RASGRP4 is the GEF that could explain the high KRAS-GTP bound levels found in Y1 cells, as predicted by the model.

**Fig. 3.**
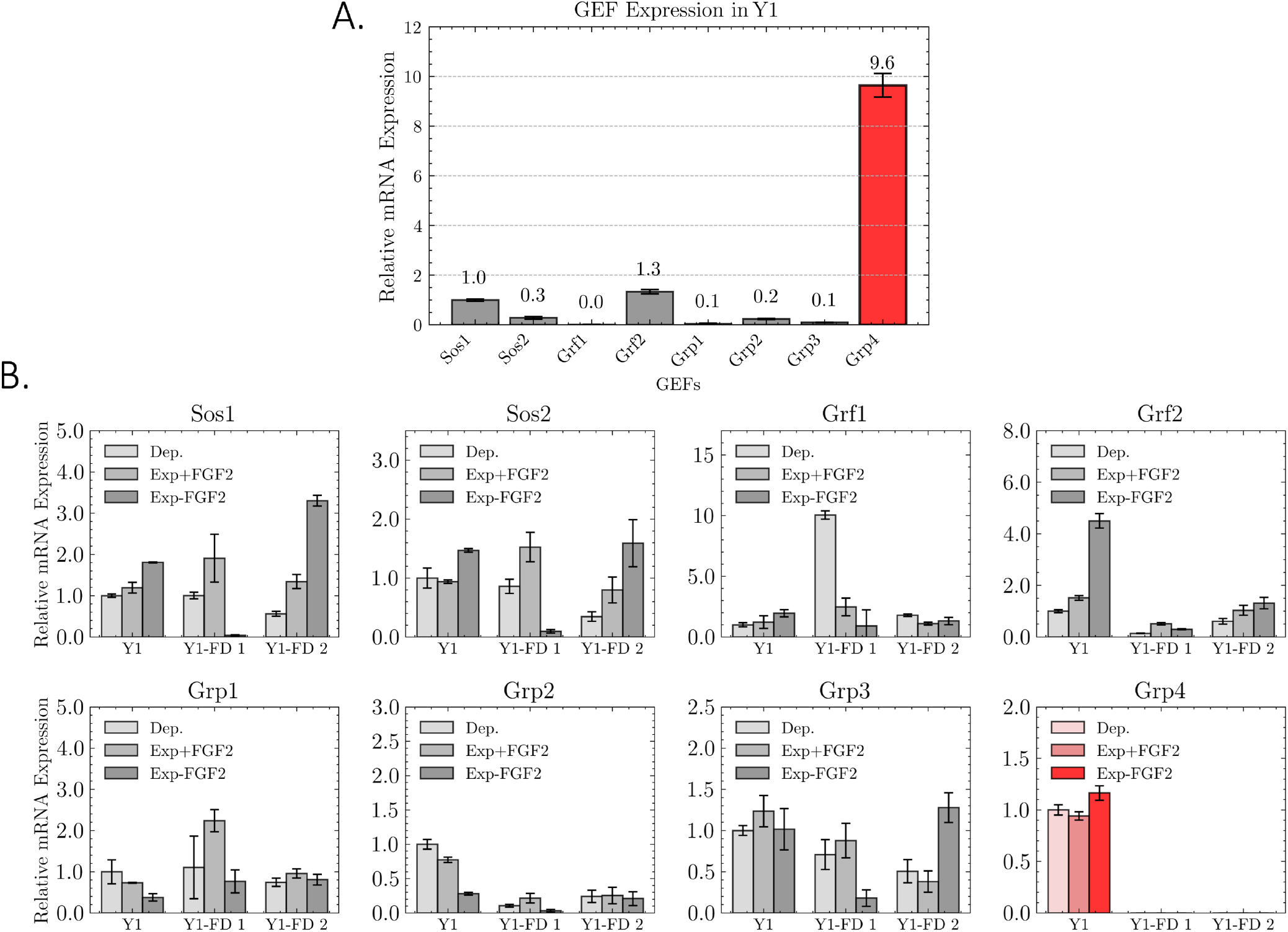
Analysis of Kras levels and expression of Ras GEFs mRNAs in parental Y1 cells and in the to Y1-FD subclones 1 and 2. **A**: mRNA expression levels of Ras GEFs in Y1 cells, normalized to *Sos1* expression. **B**: Ras GEFs mRNA levels for all cell lines, always taking as a reference the levels of the respective Ras GEF in serum deprived Y1. **Legend**: **Dep** = Deprived of FBS. **Exp+FGF2** = Cells at exponential growth treated with FGF2 **Exp-FGF2** = Cells at exponential growth with no FGF2 treatment.

### RasGrp4 depletion protects Y1 cell from cytotoxic effects caused by FGF2 due to reduced K-Ras activation

The growth curves (Figure 4A) showed that the Y1 parental cell line and ΔKRAS cell lines had slight growth difference, with both having 26.4 and 31.5 hours of doubling time, respectively. The ΔRASGRP4 and Y1-FD 1 cell line showed a high doubling time of 14.9 and 12.1 hours, respectively. The Ras activation assay (Figure 4B) has demonstrated differential Ras-GTP bound levels among the cells used in this study. Regarding Y1 cells (avg. ABS = 0.619), both ΔKRAS and ΔGRP4 cells exhibited approximately half the activation levels (avg. ABS = 0.2142 and 0.2792, respectively). In contrast, Y1-FD 1 cell displayed negligible activation (avg. ABS = 0.0354). Statistically, Tukey’s test indicated that Y1 cells were significantly different from all other groups (*p* = 2.5× 10^−7^). Furthermore, ΔGRP4 and ΔKRAS were significantly different from Y1-FD cells (*p* = 1.82× 10^−5^ and *p* = 5.75 × 10^−4^, respectively), but not significantly different from each other (*p* = 0.288). This assay corroborates with the preposition that the RASGRP4, once when removing the newly discovered GEF, it is possible to drastically change the tumor phenotype observed in Y1 cell line. Lastly, In the clonogenic assay (Figure 4C), a differential behavior between the lineages in relation to FGF2 can be observed. The ΔKRAS and Y1-FD 1 cell lines exhibited strong protection against FGF2, while GRP4 also showed protection but to a lesser extent. Additionally, the Y1-FD 1 cells can not progress if not stimulated by FGF2.

**Fig. 4.**
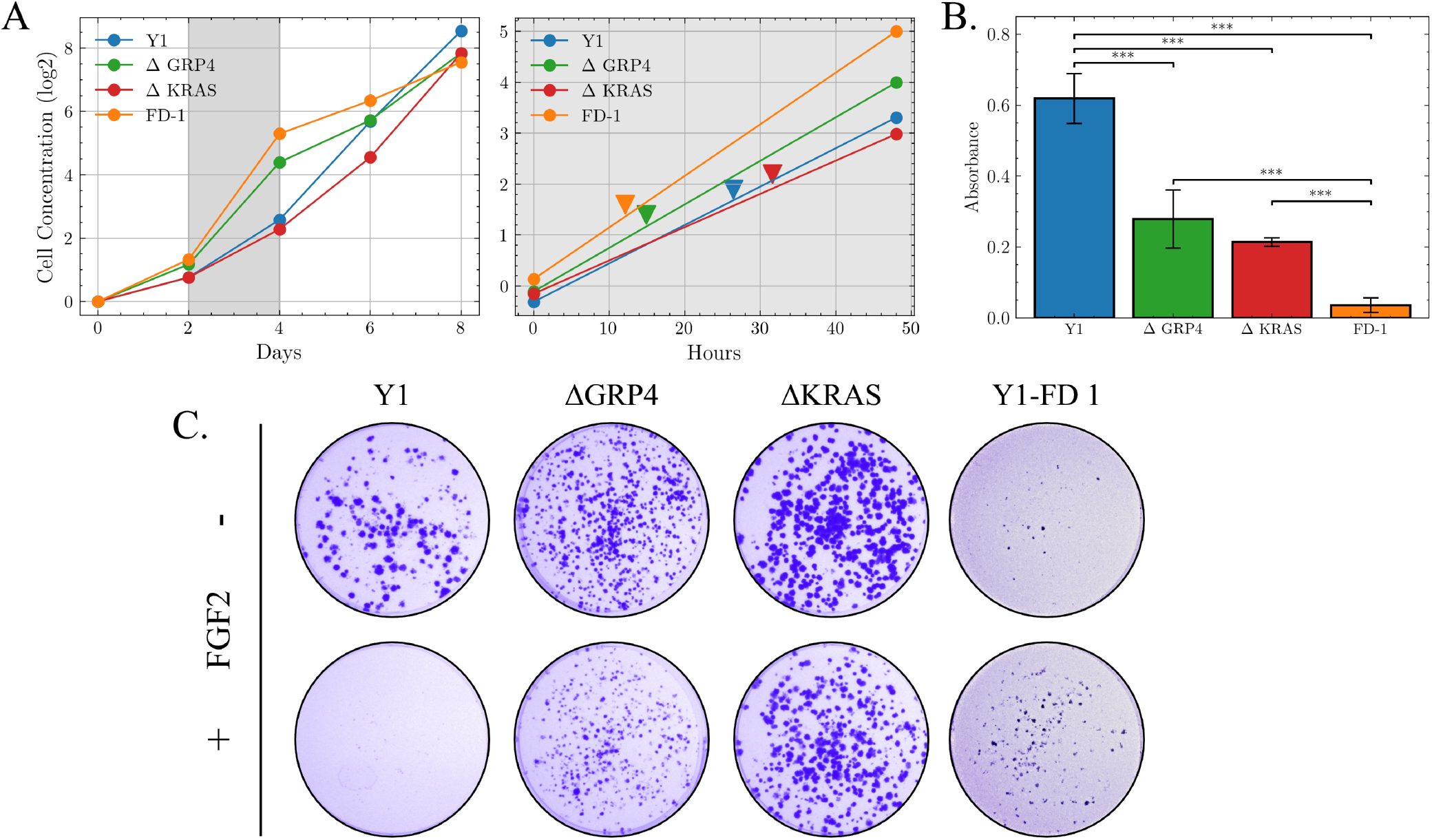
RASGRP4 depletion protects Y1 cells from cytotoxic effects caused by FGF2. **A**: Averaged cell growth curves (*n* = 3). For the assay, 0.8 *×* 10^4^ cells were plated in triplicates and treated with or without FGF2. Cells were counted over 8 days. Doubling time was calculated from the log phase (days 2–4) using log-transformed values. The individual experiments can be seen on **Supplementary Figure 1. B**: Pan-RAS detection through ELISA, comparing all the cell lines from this study. Cells were not synchronized to capture steady-state conditions. Absorbance was measured at 490 nm. Statistical analysis was done with one-way ANOVA followed by Tukey’s post hoc test. **Significance**: **\*\*\*** = *p <* 0.001. **C**: Clonogenic assay of the response to all lineages in this study versus FGF2. For this assay, 600–1000 cells per well were plated on 6-well plates, let to adhere overnight and then treated with 10 ng/mL FGF2. After that, the culture media were replaced every other day until the endpoint of 14 days.The whole experimental set can be seen on **Supplementary Figure 2. Legend**: **Y1** = Y1 Tumor cell. Δ**GRP4** = Y1 CRISPR depleted of RASGRP4 protein cell line. Δ**KRAS** = Y1 CRISPR depleted of KRAS protein cell line. **Y1-FD** = Y1 FGF2 Dependent cell line clone 1.

### RASGRP4 depletion led to lower tumor frequency

The tumor growth assay has demonstrated that the RASGRP4 depletion has changed the tumorigenic phenotype in Y1 cells. For the Y1 cell injected group, there was a rapid decline in the survival rate in Kaplan-Meier survival curves (Figure 5A) with the occurrence period ranging from the 13th and 17th day. Regarding the tumor average (Figure 5B), the Y1 group shows the most aggressive tumor progression, reaching tumors with approximately 1200 mm^3^ by the end of the observation period (60 days). In the individual profile (Figure 5C), 5/6 mice reached tumors higher than 1000 mm^3^, with the top value of 1666 mm^3^ and the lowest value of 700 mm^3^. This profile is expected for this assay, taking on account of the high-basal levels of non-mutated Kras in Y1 Cells (16–18), acting as a control for this experiment. For the ΔKRAS group, the survival rate in Kaplan-Meier survival curves (Figure 5A) show slower tumor emergence, with the occurrence period ranging from the 17th and 42nd day, with a protection rate of 1/6. When it comes to the tumor average (Figure 5B), the ΔKRAS clone has displayed a mild tumor progression, with approximately 500 mm^3^ in average by the end of the observation period. In the individual profile (Figure 5C), only 1/6 mice reached a tumor higher than 1000 mm^3^, with the top value of 1183 mm^3^ and the lowest value of 288 mm^3^. Despite the expectations for zero tumor frequency for these groups, once non-mutated Kras is responsible for the tumorigenic phenotype observed in Y1 cells, to the best of our knowledge, it is unknown the existence of other factors to contribute with that diminished tumor phenotype, such as residual mutation. Nevertheless, unveiling the adjacent mechanisms that might still be responsible for the tumor observed is not the objective of this work. As for the Y1-FD 1 line, no tumor was observed. Lastly, the ΔRASGRP4 group showed a very low decline in the survival rate in Kaplan-Meier survival curves (Figure 5A), with the occurrence period ranging from the 28th and 32nd day, with protection rate of 4/6. Regarding the tumor average (Figure 5B), the ΔRASGRP4 clone groups show low tumor progression, reaching tumors with approximately 300 mm^3^ by the end of the observation period. In the individual profile (Figure 5C), only 1 out of 6 mice reached a tumor higher than 1000 mm^3^, with the top value of 1183 mm^3^ and the lowest value of 256 mm^3^. Each individual tumor of this assay is shown in Figure 5D.

**Fig. 5.**
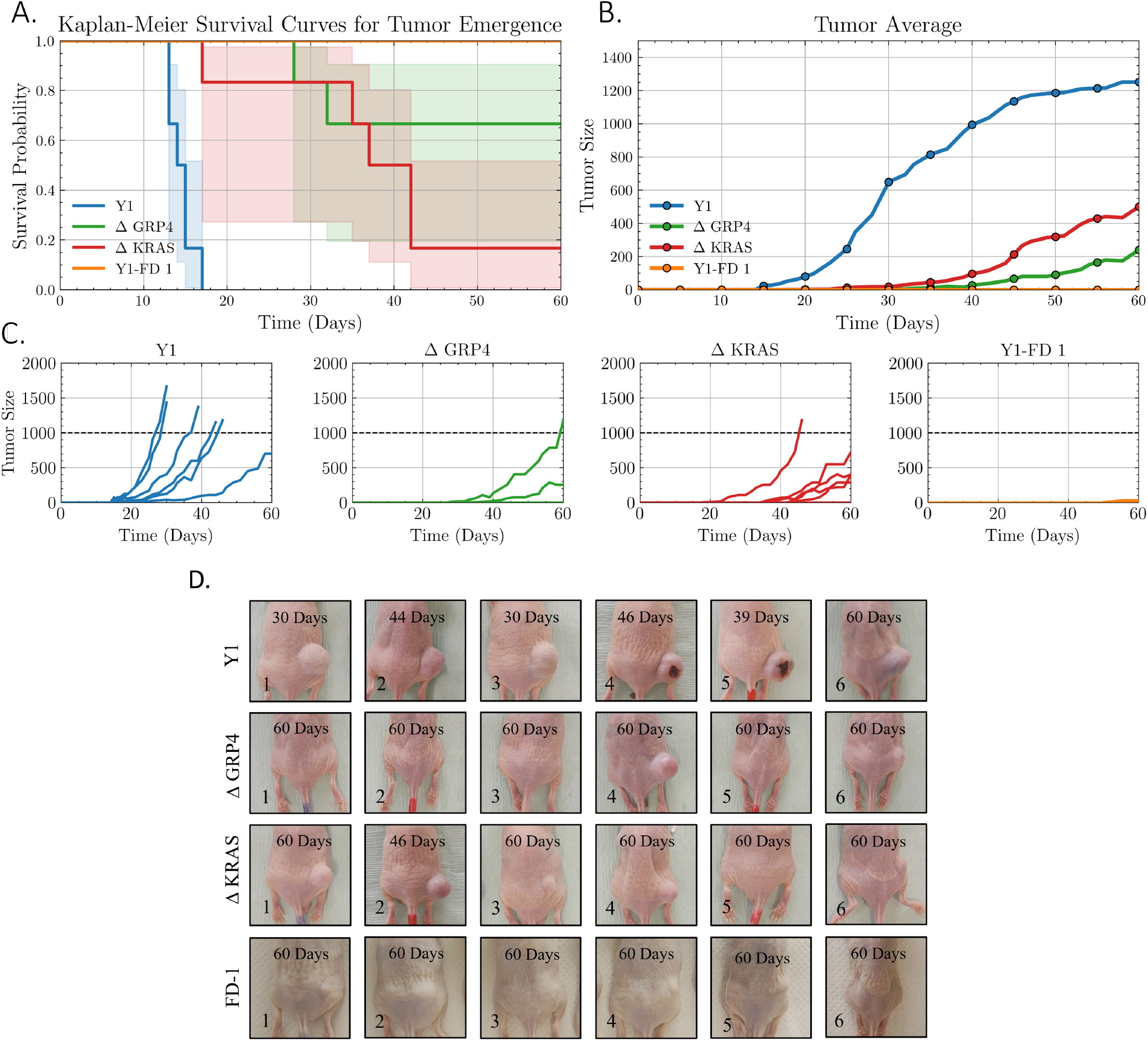
The RASGRP4 depletion led to lower tumor frequency. Six Balb/c-NUDE mice (5–8 weeks old) per group were subcutaneously inoculated with 5 *×*10^4^ cells (Y1, Y1 ΔGRP4, Y1 ΔK-RAS, or Y1-FD) in the right flank. Tumor growth and animal weight were monitored every two days. In mice with visible tumors, size was measured every other day. Tumor volume was calculated as *V* = (*L × W × W*)*/*2. Endpoint was defined by 1000 mm^3^ tumor size in controls or 60 days of observation. **A**: Kaplan-Meier survival curves for all lineages in this study. N=6. **B**: Averaged tumor growth. N=6. **C**: Individual tumor growth profile of mice during the observation period. **D**: Tumor images of mice. Each image shows the day the photo was taken, just before euthanasia when the mouse reached the experimental endpoint.The mouse weight during the observation period can be seen on **Supplementary Figure 3.Legend**: **Y1** = Y1 Tumor cell injected mouse. Δ**GRP4** = Mouse injected with Y1 CRISPR depleted of RASGRP4 protein cell line. Δ**KRAS** = Mouse injected with Y1 CRISPR depleted of KRAS protein cell line. **Y1-FD 1** = Mouse injected with Y1 FGF2 Dependent cell line clone 1.

## Discussion

In this article, we report RASGRP4 protein as a necessary factor alongside SOS in KRAS activation in murine Y1 adrenocortical tumor cells. The ODE system reproducing KRAS activation mediated by SOS resulted in mismatch with the literature (Figure1A and 2A). This prompted us to hypothesize that SOS alone could not be responsible for explaining the KRAS-GTP levels and another GEF would be needed to stabilize the system. Once accomplished, the KRAS-GTP levels matched the expected levels in the model (Figure 1B and 2B). Aiming for the missing GEF, we verified the RASGEFs profile of mRNA of the Y1 cell and Y1 FGF2 Dependent cell lines through the qPCR technique. The outcome was that the RASGRP4 was the most expressed in the Y1 cell line (Figure 3A), whereas non-existent in the Y1-FD 1 cell line (Figure 3B), leading to the hypothesis that RASGRP4 is the missing GEF in the KRAS activation model. To validate this hypothesis, we compared the ΔRASGRP4 cell line versus the Y1 cell line, which has high expression and activation of KRAS, the ΔKRAS cell line, which has the KRAS depleted and the Y1-FD cell lines, which has no RASGRP4 expression and 50% less KRAS copies (19). The cell growth curve (Figure 4A) shows that the CRISPR depleted cell lines and Y1-FD cell lines are viable and capable of growing, and in some cases as ΔGRP4 and Y1-FD even surpassing the growing rate of the Y1 cell. It is worth mentioning that the Y1-FD 1 cell line displays a more canonical growth curve: After the 4th day (after the identified log phase), the growth rate declines and ends up reaching a plateau. Differently, the other cell lines continue to show growing tendency at the 8th day, proliferating even after reaching confluence. Moreover, the Y1-FD 1 cell line is also under constant FGF2 stimulation, which we believe to contribute to the high growing rate, showing expected behavior of a normal cell lineage. The RAS-GTP ELISA assay showed that the ΔRASGRP4 cells have lower basal levels of RAS activation on their steady state in comparison to the Y1 cell (Figure 3B), corroborating with the aforementioned ODE modeling results. Regarding the response to FGF2 (Figure 3C), the ΔRASGRP4 cells showed high protection against the aforementioned deleterious effect of the FGF2 versus the Y1 cell, displaying similar results as the ΔKRAS and Y1-FD cell lines. Lastly, the tumor growth assays in Balb/c-NUDE mice with the Y1 cell versus ΔRASGRP4 cells showed reduced tumor growth and frequency for the ΔRASGRP4 cells, displaying lower tumor frequency and size even when comparing to the ΔKRAS cell line (Figure 5C). These results confirm the proposed modeling that shows that SOS cannot solely explain the KRAS-GTP levels found in the Y1 cell (Figure 1B) and RASGRP4 is a newly reported GEF that fills this system, being fundamental for the tumoral phenotype in this cell line, even not being part of the main axis of activation described for this pathway. RASGRP4 is a member of the RasGRP family (20). They are composed by a Ras exchange motif, a CDC25 homology domain, and differently than SOS, they possess a diacylglycerol-binding C1 domain and calcium-binding EF hands (20, 21). In terms of signaling, the recruitment of RASGRP4 takes a different path, activating Ras and MAPK signaling via its C1 domain in response to the diacylglycerol (DAG) (22). Their importance is reinforced in other researches, such as the work of Suiré *et al*., 2012, in which shows that RASGRP4 inhibition impairs the RAS activation in neutrophils in response to the GPCR agonist fMLP, and no other GEF, including SOS, can compensate this absence (23). Moreover, the work of Zhu *et al*., 2019, showed that the blockage of RASGRP4 considerably reduces diffuse large B cell lymphoma growth (24). Taken altogether, our findings shows that RASGRP4 is an newly important GEF for the tumorigenicity in Y1 cells, where its absence reflects in a negative tumor phenotype. Due to the conformational differences from the SOS, we speculate that this also points strongly for the influence role of the diacylglycerol regulated cell signaling pathways concomitantly with MAPK pathway to drive the tumor phenotype. These findings highlight the importance of studying GEFs, which are often overlooked, and suggest that their roles may be more complex than initially perceived. As next step, we plan to apply this approach in clinically relevant cell models, also mediated by the GEFs and GAPs. We believe that the expansion of the knowledge on the GEFs and GAPs can be game changing in cancer research as a whole.

## Methods

### Ordinary Differential Equation Modeling of K-RAS Activation Mediated by SOS in Y1 Cells

To construct the dynamic model of KRAS activation in Y1 cells, we adapted the species and reactions described by Das *et al*. (2009) (7), which incorporate the allosteric regulation of SOS by both RAS-GDP and RAS-GTP, forming a positive feedback loop. We implemented this reaction scheme as a system of ordinary differential equations (ODEs), with rate constants based on experimentally measured values. The model was then used to simulate KRAS-GTP levels under varying total concentrations of KRAS and SOS, including conditions mimicking the elevated KRAS-GTP basal levels observed in Y1 cells. To determine the steady state of the system of ODEs, all time derivatives are set to zero, reducing the model to a system of algebraic equations whose solutions represent the fixed points. The stability of each steady state is then assessed by computing the Jacobian matrix at the fixed point and analyzing its eigenvalues. If all eigenvalues have negative real parts, the steady state is locally stable, as small perturbations will decay over time.

#### Cell Lines

The Y1 murine adrenocortical carcinoma cell line was obtained from ATCC. The CRISPR-Cas9 cell lineages were derived from the parental Y1 cell line: The Y1 ΔKRAS cell line was obtained in the work of Dias *et al*. (25), while the Y1 ΔRASGRP4 was obtained during the development of this research. As for the FGF2 Dependent Y1 cell lines (Y1-FD), the clones were selected in our lab, after continuous Fibroblast Growth Factor 2 stimulation in the culture media. More details in the *FGF2 Dependent Y1 Cells Obtaining* section.

#### Cell Culture

The Y1 parental and CRISPR depleted cell lines were grown at 37°C in 5% CO2 atmosphere in Dulbecco’s modified Eagle’s Medium (DMEM; # 11885084, Gibco, USA) supplemented with 10% Fetal Bovine Serum (FBS; #S0011, Vitrocell, Brazil). For Y1-FD cell lines, 10 ng/ mL FGF2 was added to the media. Before the execution of all experiments, the synchronization in the phase G0/G1 by serum starvation was performed by the removal of the FBS supplemented media, PBS A (#14190-136, Gibco, USA) wash and FBS-Free DMEM addition for 48h prior to any stimulation.

#### RNA Extraction

The total RNA from cells was extracted with the Illustra RNAspin Mini RNA Isolation kit (#25-0500-72; GE Healthcare) following the manufacturer’s instructions. In brief, the samples were treated with DNAse I, and Immediately after extraction, the RNA fractions were quantified in the spectrophotometer Nanodrop (Nanodrop, Thermo Fisher Scientific; Waltham, Massachusetts, EUA) at wavelengths of 280 nm and 260 nm and stored at −80ºC until use. Next, the complementary DNA (cDNA) was synthesized by reverse transcription with the SuperScriptTM III Reverse Transcriptase kit (#18080085; Thermo Fisher Scientific, USA) and used as a template for the real-time PCR reaction.

#### PCR

To experimentally test the aforementioned computational prediction, we investigated the expression levels of members of the known Ras-GEFs families (Sos, Grp and Grf) by RT-qPCR Primers designed using the Primer-BLAST tool (26) (Table 1), according to cDNA consensus sequences deposited in GenBank (27), with specificity checked by performing an *in-silico* PCR using the UCSC tools (28). The experimental conditions were: (1) Deprived of FBS for 24h, (2) exponential growth - FGF2 for 24h and (3) exponential growth + FGF2 10 ng/ mL for 24h, for both in Y1 parental and Y1-FD cells. The reaction was performed by using the SYBR® GREEN PCR Master Mix kit (#4309155; ThermoFisher Scientific) with 40 ng of RT RNA and 2.5 µL 300 nM of forward and reverse primer oligonucleotide solutions, following the manufacturer’s instructions. Amplifications were performed in cycles of denaturation reactions (95°C, 15 sec), annealing and polymerization (60°C, 60 sec) in a thermocycler (StepOne Plus, Applied Biosystems, USA), using the own StepOne software for data collection. For each sample, the efficiency of the primers was determined by the software LinRegPCR tools (LinRegPCR v. 12.17 software) (29), which was used for relative quantification of the data obtained by the Pfaffl method (30). All the results were quantified relatively, taking the untreated condition as a comparison. Hprt was used as an endogenous normalizing control for the mRNA expression data.

**Table 1.**
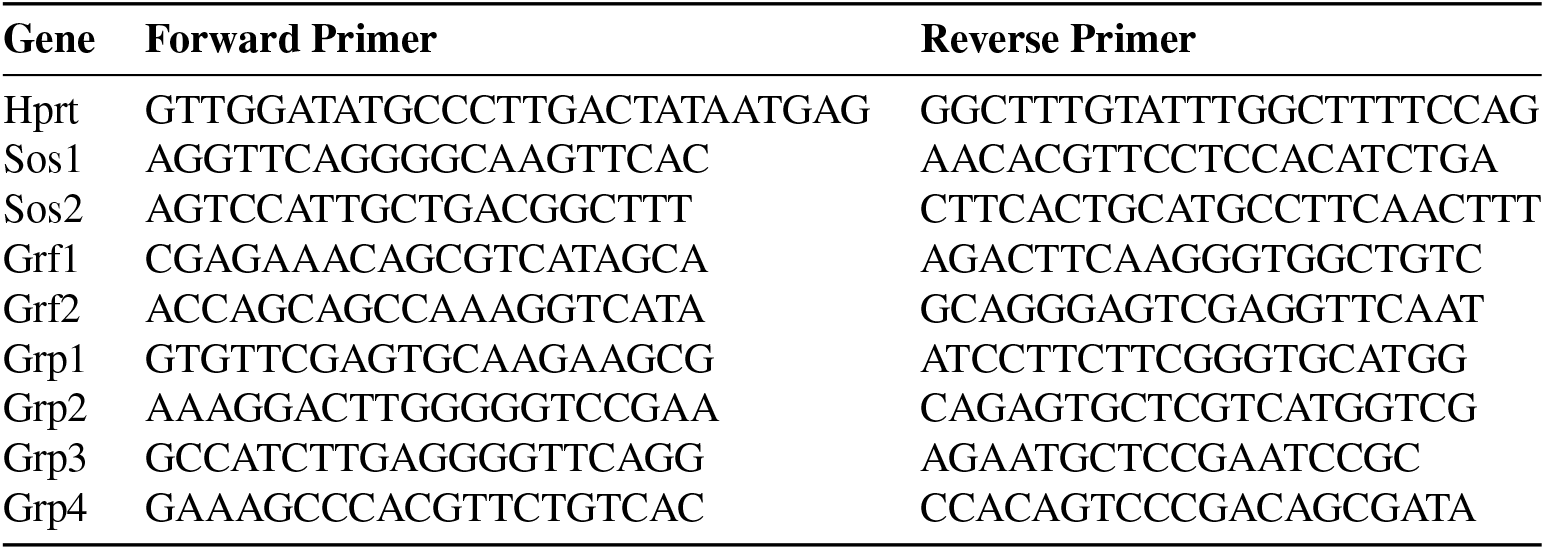
Sequences of primers used for gene amplification by PCR.

### Y1 ΔRASGRP4 and Y1 ΔKRAS cell lines obtaining thought CRISPR-Cas9

For this research project, we used the same Y1 cell lines depleted of the Kras gene which was used in the publication Dias *et al*. 2019 (25) and Y1 recently depleted from the Rasgrp4 gene in our laboratory, with both obtained with similar methods. Briefly, we developed five sgRNAs for the respective genes using the CRISPR design tool, from MIT (*https://www.ensembl.org/index.html*; *https://crispor.gi.ucsc.edu/* (31), which can be seen in the Table 2. The oligos were cloned into a LentiGuide-Puro plasmid (#52963, Addgene, USA), using the methodology described by Sanjana et al. (32). For the lentivirus production, the LentiGuide-Puro constructs, psPAX2 (#12260, Addgene, USA) and pCMV-VSV-G (#8454; Addgene, USA) were transfected into HEK293T cells using lipofectamine 3000 reagent (#L3000015, ThermoFisher Scientific, USA), following the manufacturer’s recommendations. After 48h of transfection, the viral supernatants were collected and filtered. The Y1 cells were then cotransduced with the constructs LentiCas9-Blast and LentiGuide-Puro 8 µg/ mL hexadimethrin bromide (#sc-134220, Santa Cruz Biotechnology, USA). After 48h of transduction, Y1 cells were selected with puromycin 3 µg/ mL and blasticidin 7 µg/ mL for 7 days. Finally, these cells had several clones selected and subcultured, where they were finally tested using the western blot technique with antibodies against the proteins in which their genes were knocked out. Lastly, to obtain individual cells for colony formation, we calculated a serial dilution to distribute 50 cells in three technical replicates across 96-well microplates. Over time, the cells were subcultured sequentially from 96-well plates to 24-well, 12-well, and finally 6-well plates. Once colonies formed, they were transferred to culture flasks for expansion or cryopreserved in a freezing solution (40% FBS, 50% DMEM, and 10% DMSO) at −80°C.

**Table 2.**
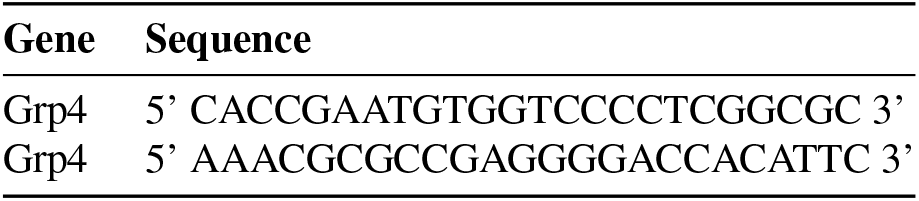
sgRNA sequences designed for CRISPR targeting of RasGrp4 gene.

**Table 3.** Kinetic parameters used in this study for modeling Ras activation dynamics in Y1 cells. Rate constants were assigned standard biochemical units based on reaction order. Bimolecular association rates (e.g., *k*_1_, *k*_2_) are expressed in *µ*M^−1^ · s^−1^, while unimolecular dissociation rates (*d*_1_, *d*_2_) and catalytic rates (*k*_cat_) are in s^−1^. Michaelis constants (*K*_*m*_) are reported in *µ*M, and maximum velocities (e.g., *V*_5_) in *µ*M · s^−1^. These conventions follow classical mass-action and Michaelis–Menten kinetics, as adopted in previous modeling efforts (7).

| Parameter | Description | Value (Y1 cells) |
| --- | --- | --- |
| $k_1$ | SOS binding Ras-GDP (kon) | $1.8 \times 10^{-4} \mu\text{M}^{-1} \cdot \text{s}^{-1}$ |
| $d_1$ | SOS–Ras-GDP dissociation (koff) | $3.0 \text{ s}^{-1}$ |
| $k_2$ | SOS binding Ras-GTP (kon) | $1.7 \times 10^{-4} \mu\text{M}^{-1} \cdot \text{s}^{-1}$ |
| $d_2$ | SOS–Ras-GTP dissociation (koff) | $0.04 \text{ s}^{-1}$ |
| $k_{3cat}$ | Catalysis with Ras-GTP (allosteric site) | $0.038 \text{ s}^{-1}$ |
| $K_{3m}$ | Michaelis constant for $k_{3cat}$ | $1640 \mu\text{M}$ |
| $k_{4cat}$ | Catalysis with Ras-GDP (allosteric site) | $0.003 \text{ s}^{-1}$ |
| $K_{4m}$ | Michaelis constant for $k_{4cat}$ | $9120 \mu\text{M}$ |
| $V_5$ | RasGAP-mediated deactivation | $0.1 \times [\text{GAP}] \mu\text{M} \cdot \text{s}^{-1}$ approx. |
| $K_{5m}$ | Michaelis constant for GAP | $107 \mu\text{M}$ |
| $k_{6cat}$ | Additional GEF-mediated exchange | $0.01 \text{ s}^{-1}$ |
| $K_{6m}$ | Michaelis constant for additional GEF | $1836 \mu\text{M}$ |

### FGF2 Dependent Y1 Cells Obtaining

The cell lines used in this study were kindly provided by Dr. Tatiana Guimarães de Freitas Matos. Y1 cells were plated at a density of 6.4 × 10^2^ cells/cm^2^ in triplicates. After plating, the cells were kept at rest for 24 hours before receiving treatment with FGF2 at a concentration of 10 ng/mL. The treatment was maintained with the selection medium containing FGF2 being renewed three times a week. In the first 24 hours of exposure to FGF2, intense cell death was observed, which continued in the subsequent days. After 15 to 20 days of treatment, the emergent colonies were isolated using cloning rings, with the cells being trypsinized and transferred to 12-well plates, where each well contained cells from a single colony. The isolated cells were expanded in the presence of FGF2. The clones obtained were called Y1-FD (FGF2 Dependent). For this study, we worked with two subclones: Y1 subclone 1 (Y1-FD 1) and Y1 subclone 2 (Y1-FD 2). Y1-FD 1 was used for all the experimentation described, whereas Y1-FD 2 was used only for PCR analysis.

### Ras-GTP detection with G-LISA Assay

The colorimetric assay was performed in quintuplicate using RAS G-LISA Activation Assay Kit (#BK131, Cytoskeleton, USA) according to the manufacturer’s recommendations and read in a spectrophotometer by measuring absorbance at 490 nm. Differently than the other assays, the cells were now synchronized with the objective of capturing their steady state. Statistical differences between groups were assessed using one-way analysis of variance (ANOVA), followed by Tukey’s post hoc test for multiple comparisons. The analysis was performed using Python with the statsmodels (33) and scikit-posthocs (34) libraries.

### Clonogenic and Viability Assays

For this assay, 600-1000 cells per well were plated on 6-well plates, let to adhere overnight and then treated with 10 ng/mL FGF2. After that, the culture media were replaced every other day until the endpoint of 14 days. Cells were then washed with PBS, fixed, and stained in a fixing/staining solution (0.05% crystal violet, 4% formaldehyde, 10% methanol in PBS) and washed abundantly. Images were acquired using UVITEC Cambridge (UVITEC, UK).

### Growth Curves

For this assay, 0.8 × 10^4^ cells were plated on 12-well plates in triplicates in 3 independent experiments, let to adhere overnight. After that, the cells were synchronized as described previously. The cells were then treated with or without FGF2 in DMEM 10% FBS. In the 8 days, cells were harvested and counted in a Z2 Beckman Coulter® (Beckman, USA) counter. The log phase of cell growth was identified between days 2 and 4. Doubling time was calculated using cell counts from this interval (48 h). Values were log-transformed, and doubling time was determined using the formula:

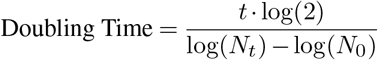

where *t* is the time interval (48 h), *N*_0_ is the cell number at day 2, and *N*_*t*_ is the cell number at day 4.

### Mouse Tumor Assay

For all experiments 6 Balb/c-NUDE mice, 5-8 weeks old, per group were used. The mice were injected with 5×10^4^ tumor cells Y1, Y1 ΔGRP4, Y1 ΔK-RAS and FGF2 Dependent Cell lines (Y1-FD), subcutaneously, on the back of the right flank. All groups were examined at each 2 days, animal by animal, measuring weight and tumor presence. In positive cases (tumor visible and palpable), tumors were measured for length and width, every other day. The tumor volume is calculated by the formula V = (L x W x W)/2, where V is tumor volume, W is tumor width, and L is tumor length. The endpoint was done when the control animals reach tumors with 1000mm^3^ or 60 days of assay. The data was processed with Pandas (35) 2.2.3 libraries for calculations, KaplanMeierFitter from Lifelines 0.30.0 (36). Experiments using mice were carried out after approval of the Committee on Ethics in the Use of Animals of the Butantan Institute with the number CEUAIB 3976280723.

### Visualization

For generating the figures, we used Matplotlib 3.9.0 (37) for plotting and Science Plots (38) for styling.

## Acknowledgements

- Butantan Institute
- The São Paulo Research Foundation (FAPESP) Grants:
  - #13/07467-1
  - #19/21619-5
  - #19/24580-2
  - #20/08555-5
  - #20/10329-3
  - #21/04355-4

- Brazilian Federal Agency for Support and Evaluation of Graduate Education (CAPES) Grant: #001

## Data Availability

The ODE model script and data used in this paper can be found in the @anthraxodus/Montoni2025_RASGRP4_KeyGEF_Y1 GitHub repository.

---

**Synopsis:**
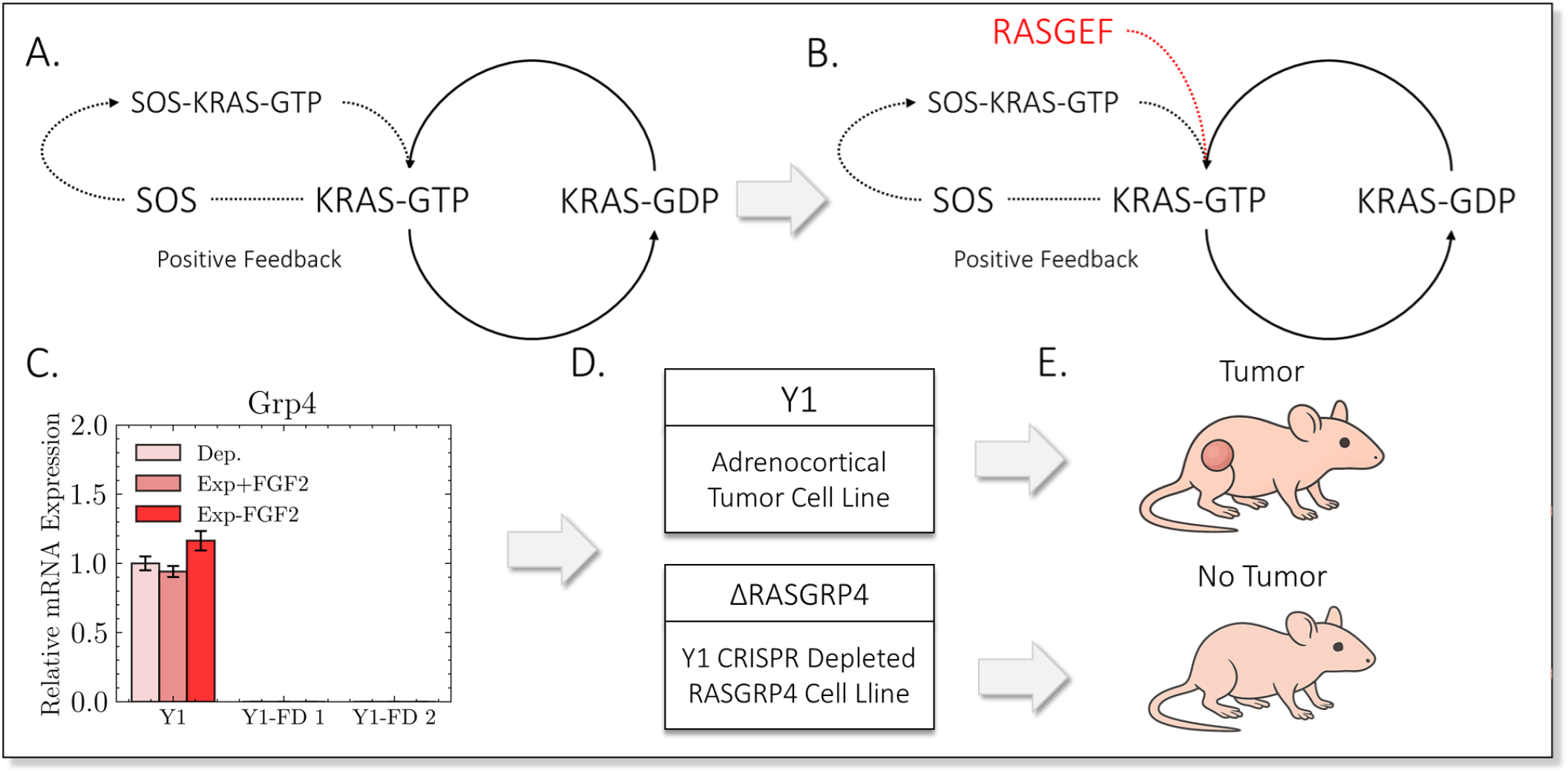
**A**: Initial ODE modeling of KRAS activation mediated by SOS. The levels of activation in the steady state of the model did not match the expected levels. **B**: After adding another GEF, the levels matched, pointing that SOS was not solely responsible for the activation. **C**: Further PCR investigation revealed Rasgrp4 to be non-existent in Y1-FD Cells, suggesting to be the missing GEF in the system. **D**: A CRISPR against RASGRP4 was performed. **E**: The mouse tumor assay comparing Y1 and Y1 ΔRASGRP4 cells showed that depleted RASGRP4 Y1 cells can not progress in tumorigenesis, corroborating with the hypothesis that SOS cannot solely activate KRAS in Y1 cells.

## Supplementary Information

**Supplementary Figure 1:**
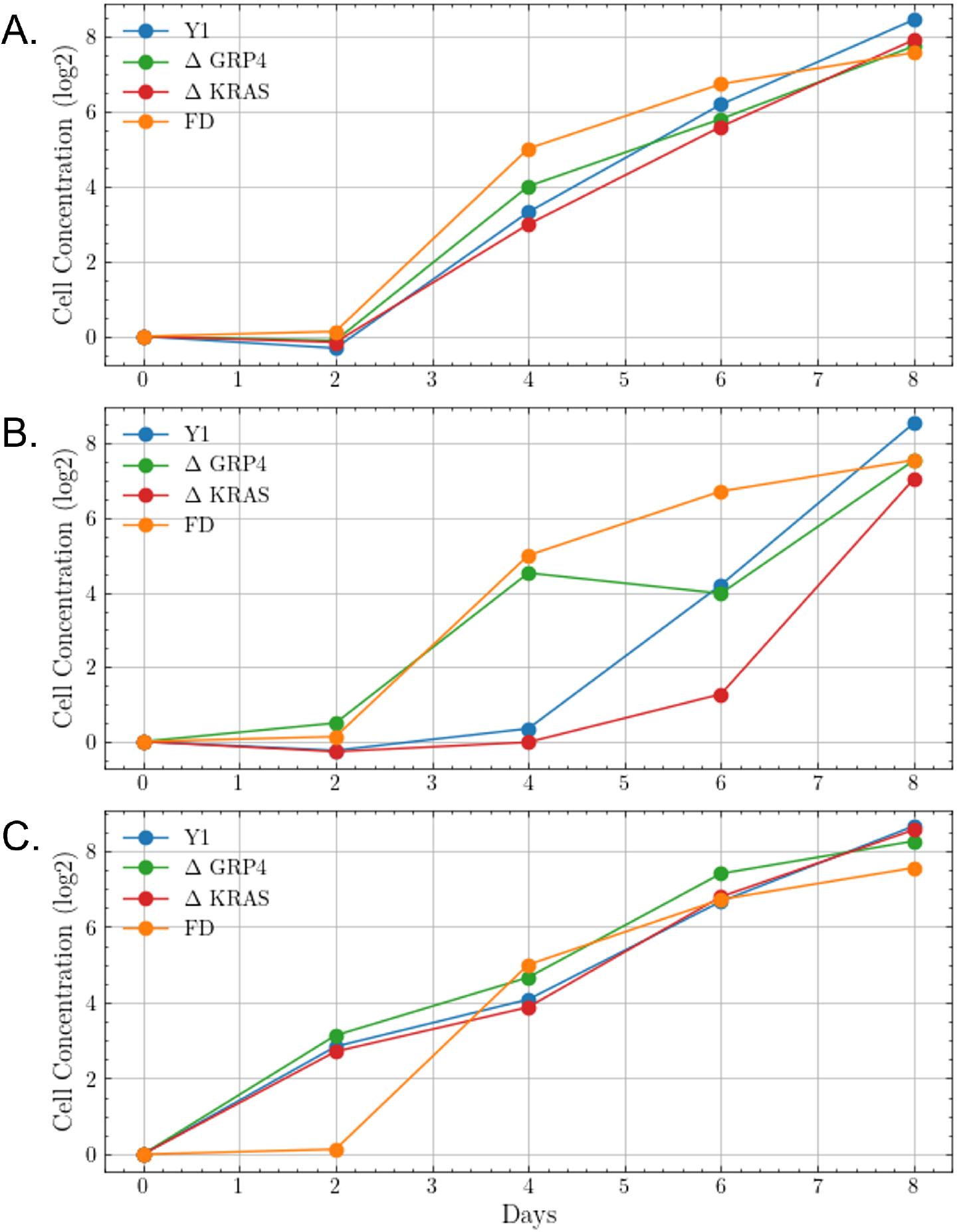
Individual profile of the growth curves. **A**: Experimental replicate 1. **B**: Experimental replicate 2. **C**: Experimental replicate 3. Legend: **Y1** = Y1 Tumor cell. Δ**GRP4** = Y1 CRISPR depleted of RASGRP4 protein cell line. Δ**KRAS** = Y1 CRISPR depleted of KRAS protein cell line. **Y1-FD** = Y1 FGF2 Dependent cell line clone 1.

**Supplementary Figure 2:**
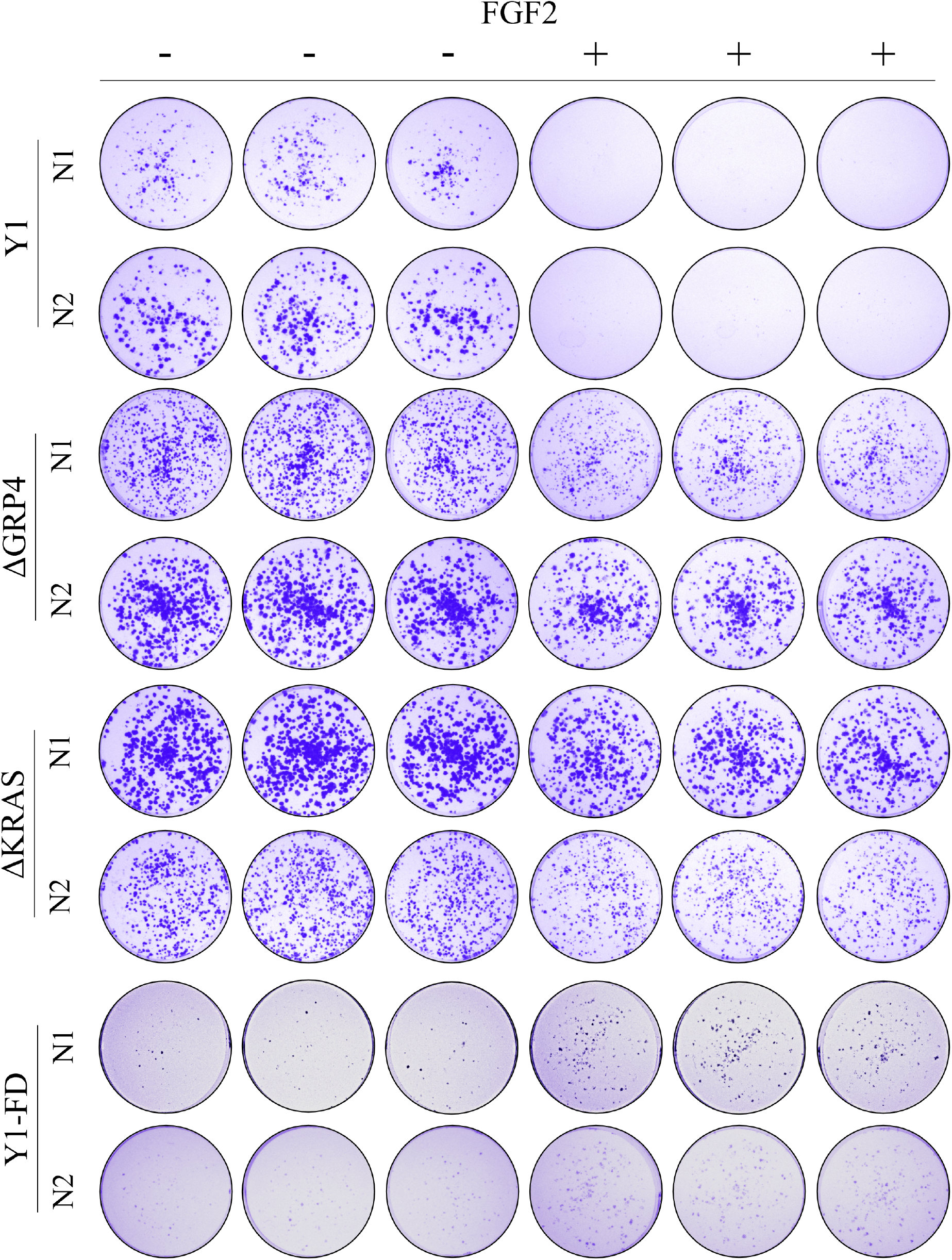
Clonogenic assay of the response to all lineages in this study versus FGF2. For this assay, 600–1000 cells per well were plated on 6-well plates, let to adhere overnight and then treated with 10 ng/mL FGF2. After that, the culture media were replaced every other day until the endpoint of 14 days. **Legend**: **Y1** = Y1 Tumor cell. Δ**GRP4** = Y1 CRISPR depleted of RASGRP4 protein cell line. Δ**KRAS** = Y1 CRISPR depleted of KRAS protein cell line. **Y1-FD** = Y1 FGF2 Dependent cell line.

**Supplementary Figure 3:**
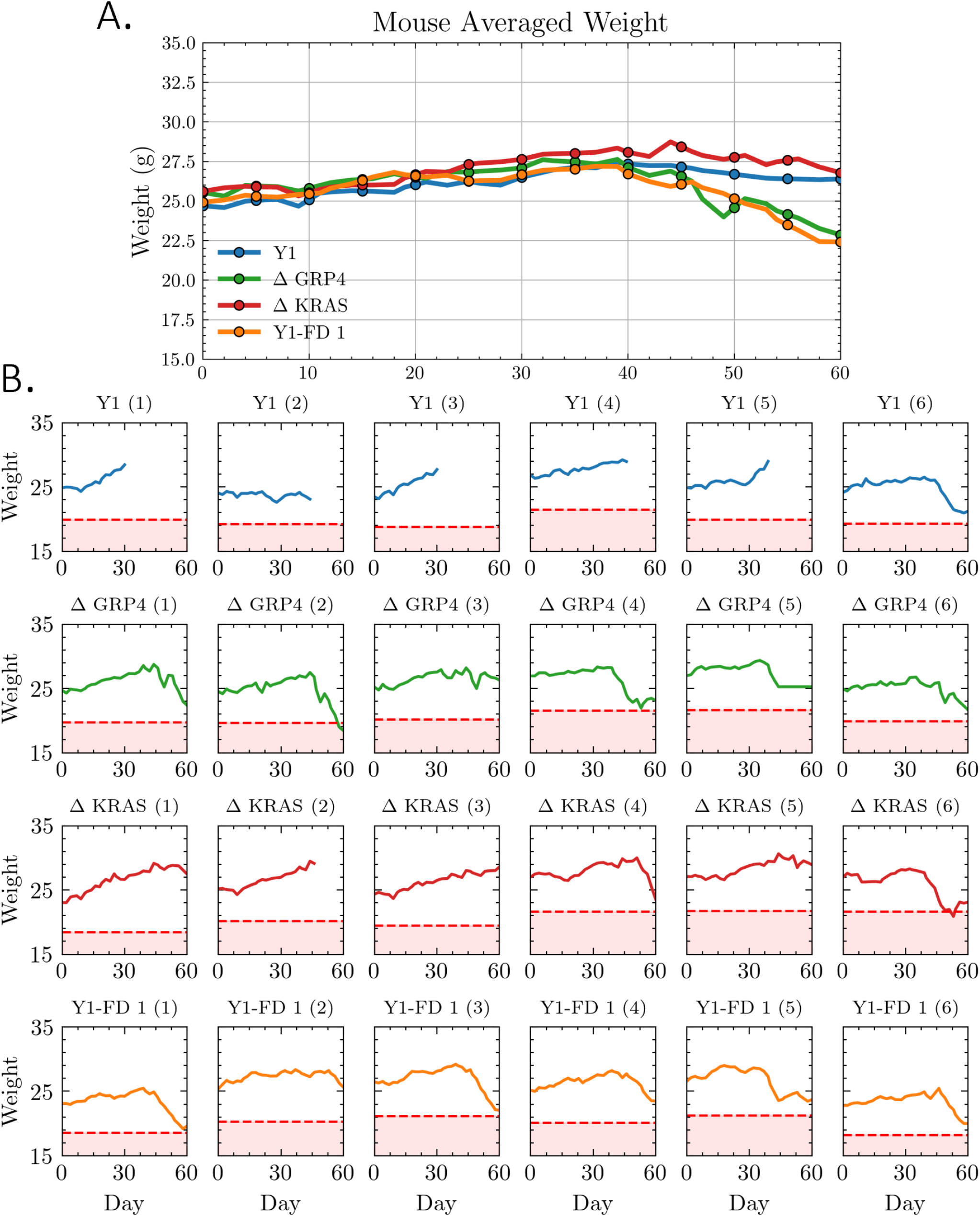
Mouse weight during the tumor assay observation days. **A**: Averaged weight by group (N=6). **B**: Individual mouse weight during the observation days.The red dashed line represents the rate of 20% of weight lost, which is calculated individually. Legend: **Y1**: Y1 Tumor cell transfected mouse. Δ**GRP4**: Mouse transfected with Y1 CRISPR depleted of RASGRP4 protein cell line. Δ**KRAS**: Mouse transfected with Y1 CRISPR depleted of KRAS protein cell line. **Y1-FD**: Mouse transfected with Y1 FGF2 Dependent cell line clone 1.

