## Supplementary material for "RASGRP4 is a key factor in KRAS activation mediated by SOS in Y1 mouse tumor cell line": Source Tables and Figures: graph_generator.py

```
"""

graph_generator.py.

Module that contains classes for plotting of yielded results.

    Copyright (c) 2019. Marcelo S. Reis.

    This module is free software: you can redistribute it and/or modify it
    under the terms of the GNU General Public License as published by the
    Free Software Foundation, either version 3 of the License, or (at your
    option) any later version.
    ReDyMo is distributed in the hope that it will be useful, but WITHOUT
    ANY WARRANTY; without even the implied warranty of MERCHANTABILITY or
    FITNESS FOR A PARTICULAR PURPOSE. See the GNU General Public License
    for more details.
    You should have received a copy of the GNU General Public License along
    with ReDyMo. If not, see <http://www.gnu.org/licenses/>.

"""
import math
import matplotlib.pyplot as plt
from mpl_toolkits.mplot3d import Axes3D

class GraphGenerator:

  # In this constructor, we define the chemical 
  # species and also the output file name and path.
  # We also define if the y-axis of graphs will be
  # at absolute or logarithm scale.
  #
  def __init__(self, x, y, z, plot_file, x_lim, y_lim, z_lim):
    self.x = x
    self.y = y
    self.z = z
    self.x_lim = x_lim
    self.y_lim = y_lim
    self.z_lim = z_lim
    self.plot_file = plot_file

    self.is_log = True
    self.log_base = 100

  # This method plots a graph to depict results
  # from a numerical simulation of the ODE system.
  # (i.e., a timecourse of selected chemical species).
  #
  def plot_simulation_graph(self, Y, t , species):

    plt.plot(t, Y[:, 0], 'b', label = species[0])
    plt.plot(t, Y[:, 3], 'b', label = species[3])
    plt.plot(t, Y[:, 4], 'g', label = species[4])
    plt.legend(loc = 'best')
    plt.xlabel('time (seconds)')
    plt.ylabel('concentration (molecules/um^3)')
    plt.grid()
    plt.show()

  # This method plots a 2D steady state graph, in
  # which the y-axis may be eather in absolute or
  # logarithm scale (self.is_log defines this property).
  #
  def plot_steady_state_graph(self, point, color):
    
    fig = plt.figure()
    ax = fig.add_subplot(111)

    if self.is_log == True:
      for i in range(0,len(point[self.z])):
        point[self.z][i] = math.log(point[self.z][i]) / math.log(self.log_base)
      ax.set_xlabel(self.x + ' (concentration)')
      ax.set_ylabel(self.z + ' (log concentration)')
      self.z_lim = [-0.5, math.ceil(math.log(self.z_lim[1]) / math.log(self.log_base))]
    else:
      ax.set_xlabel(self.x + ' (concentration)')   # Ommiting units
      ax.set_ylabel(self.z + ' (concentration)')   # (molecules/um^3).

    ax.set_ylim(self.z_lim) 
    ax.set_xlim(self.x_lim)   
    ax.scatter(point[self.x], point[self.z], c=color)
    plt.grid()
    plt.show()

# End of simulator.py.
#
```
