## Supplementary material for "RASGRP4 is a key factor in KRAS activation mediated by SOS in Y1 mouse tumor cell line": Source Tables and Figures: instance_generator.py

```
"""

instance_generator.py.

Module that contains classes for generation
of Ras small GTPase dynamic model instances.

    Copyright (c) 2019. Marcelo S. Reis.

    This module is free software: you can redistribute it and/or modify it
    under the terms of the GNU General Public License as published by the
    Free Software Foundation, either version 3 of the License, or (at your
    option) any later version.
    ReDyMo is distributed in the hope that it will be useful, but WITHOUT
    ANY WARRANTY; without even the implied warranty of MERCHANTABILITY or
    FITNESS FOR A PARTICULAR PURPOSE. See the GNU General Public License
    for more details.
    You should have received a copy of the GNU General Public License along
    with ReDyMo. If not, see <http://www.gnu.org/licenses/>.

"""

import sympy as sp

class RasModel:

  def __init__(self, ras, sos, gap, gef):

    # Chemical species.
    #
    self.GAP             = gap
    self.GEF             = gef
    self.Total_Ras       = ras
    self.Total_SOS       = sos

    # Rate constants.
    #
    # (Fábio Nakano used 2 units for k3cat)
    #
    self.param = {\
                   'k1'    : 1.8e-4,\
                   'd1'    : 3.0,\
                   'k2'    : 1.7e-4,\
                   'd2'    : 0.04,\
                   'k3cat' : 0.038,\
                   'K3m'   : 1.64e3,\
                   'k4cat' : 0.003,\
                   'K4m'   : 9.12e3,\
                   'V5'    : 0.1 * self.GAP,\
                   'K5m'   : 1.07e2,\
                   'k6cat' : 0.01,\
                   'K6m'   : 1836\
                 }

  # RasGRP1: k6cat = 0.01, K6m = 1836
  #

  # Method that receives a set of initial concentrations, update
  # the concentration attributes with them and returns the ODE system
  # ready for usage by odeint numeric integrator.
  #
  def IVP(self,y,t):    # IVP = initial value problem
    
    SOS, SOS_allo_RasGDP, SOS_allo_RasGTP, RasGDP, RasGTP = y

    # Definition of ODE system.
    #
    dydt = [- (self.param['k1'] * SOS * RasGDP) + (self.param['d1'] * SOS_allo_RasGDP) \
            - (self.param['k2'] * SOS * RasGTP) + (self.param['d2'] * SOS_allo_RasGTP), \
  
            + (self.param['k1'] * SOS * RasGDP) - (self.param['d1'] * SOS_allo_RasGDP), \

            + (self.param['k2'] * SOS * RasGTP) - (self.param['d2'] * SOS_allo_RasGTP), \

            - (self.param['k1'] * SOS * RasGDP) \
            + (self.param['d1'] * SOS_allo_RasGDP) \
            - (self.param['k3cat'] * SOS_allo_RasGTP * RasGDP)/(self.param['K3m'] + RasGDP) \
            - (self.param['k4cat'] * SOS_allo_RasGDP * RasGDP)/(self.param['K4m'] + RasGDP) \
            + (self.param['V5'] * RasGTP)/(self.param['K5m'] + RasGTP) \
            - (self.param['k6cat'] * self.GEF * RasGDP)/(self.param['K6m'] + RasGDP), \

            - (self.param['k2'] * SOS * RasGTP) \
            + (self.param['d2'] * SOS_allo_RasGTP) \
            + (self.param['k3cat'] * SOS_allo_RasGTP * RasGDP)/(self.param['K3m'] + RasGDP) \
            + (self.param['k4cat'] * SOS_allo_RasGDP * RasGDP)/(self.param['K4m'] + RasGDP) \
            - (self.param['V5'] * RasGTP)/(self.param['K5m'] + RasGTP) \
            + (self.param['k6cat'] * self.GEF * RasGDP)/(self.param['K6m'] + RasGDP)]

    return dydt


  # Method that receives an initial guess and returns the yielded
  # righthanded sides of each ODE to allow the search for roots by the
  # non-linear solver.
  #
  def non_linear_equations(self, y):

    SOS, SOS_allo_RasGDP, SOS_allo_RasGTP, RasGDP, RasGTP = y
  
    eq1 = self.Total_SOS - SOS - SOS_allo_RasGTP - SOS_allo_RasGDP

    eq2 = + (self.param['k1'] * SOS * RasGDP) - (self.param['d1'] * SOS_allo_RasGDP)

    eq3 = + (self.param['k2'] * SOS * RasGTP) - (self.param['d2'] * SOS_allo_RasGTP)

    eq4 = self.Total_Ras - RasGDP - RasGTP - SOS_allo_RasGTP - SOS_allo_RasGDP

    eq5 = + (self.param['d2'] * SOS_allo_RasGTP) \
          - (self.param['k2'] * SOS * RasGTP) \
          + (self.param['k3cat'] * SOS_allo_RasGTP * RasGDP)/(self.param['K3m'] + RasGDP) \
          + (self.param['k4cat'] * SOS_allo_RasGDP * RasGDP)/(self.param['K4m'] + RasGDP) \
          - (self.param['V5'] * RasGTP)/(self.param['K5m'] + RasGTP) \
          + (self.param['k6cat'] * self.GEF * RasGDP)/(self.param['K6m'] + RasGDP)

    return (eq1, eq2, eq3, eq4, eq5)


# End of instance_generator.py.
#
```
