## Supplementary material for "RASGRP4 is a key factor in KRAS activation mediated by SOS in Y1 mouse tumor cell line": Source Tables and Figures: main.py

```
"""

Steady-state-explorer main program.

    Copyright (c) 2019. Marcelo S. Reis.

    This module is free software: you can redistribute it and/or modify it
    under the terms of the GNU General Public License as published by the
    Free Software Foundation, either version 3 of the License, or (at your
    option) any later version.
    ReDyMo is distributed in the hope that it will be useful, but WITHOUT
    ANY WARRANTY; without even the implied warranty of MERCHANTABILITY or
    FITNESS FOR A PARTICULAR PURPOSE. See the GNU General Public License
    for more details.
    You should have received a copy of the GNU General Public License along
    with ReDyMo. If not, see <http://www.gnu.org/licenses/>.

"""

import numpy as np

from graph_generator import *

from instance_generator import *

from robustness import *

from steady_state import *

from simulator import *

import sys


def main():

  # Definition of output files.
  #
  dat_file = 'output/steady_state_data.dat'
  png_file = 'output/steady_state_graph.png'

  # Instantiation of a model simulator.
  #
  simulator = Simulator()

  # Instantiation of a robustness procedure employing a 
  # 10-fold sensitivity analysis.
  #
  robustness = Robustness(10)

  # Instantiation of a steady state evaluator.
  #
  evaluator = SteadyState('SOS,SOS_allo_RasGTP,SOS_allo_RasGDP,RasGDP,RasGTP')

  # We set the ranges of initial concentrations; that is, we define 
  # the region of the state space that will be explored in this search.
  # 
  # Since we will start to explore, for any species, from a value close
  # to zero (or zero, in the case of [GEF]), we need to store here just
  # the maximum concentration value for each species.
  #
  # Ras and SOS here stand for the total concentrations of those species.  SOS = 4000
  #
  max_concentration = \
         {'Ras' : 20000,\
          'SOS' : 1000,\
          'GAP' : 125,\
          'GEF' : 1000} 

  color = [] 
  weak  = []
  point = {'[SOS]' : [], '[RasGTP]' : [], '[GAP]' : []}

  ras = max_concentration['Ras']
  gef = max_concentration['GEF']  
  gap = max_concentration['GAP']  # 125 was used in the Das' paper.

  # Definition of the number of seconds of a numeric simulation. 
  #
  simulation_time = 1000

  # Instantiation of a graph plotter.
  #
  plotter = GraphGenerator('[SOS]','[GAP]','[RasGTP]', png_file,\
    [0, max_concentration['SOS']], [], [0,max_concentration['Ras']])

  for sos in range(10, max_concentration['SOS'], 10):

    # Instantiation of a Ras small GTPase dynamic model.
    #
    model = RasModel(ras, sos, gap, gef)

    # Definition of simulation time frame (0 to 1000 seconds).
    #
    t = np.linspace (0, simulation_time, 1000001)
       
    # Array with initial concentrations.
    #
    y0 = (sos, 0, 0, ras, 0)

    # Simulation of the Ras GTPase model.
    #
    Y = simulator.simulate(model, y0, t)

    # Plotting simulation results (uncomment here for debug purposes).
    #
    #plotter.plot_simulation_graph(Y, t, ('SOS','','','RasGDP','RasGTP'))
    #sys.exit()

    simulation_guess = Y[-1,:] # guess for the steady state solver.

    # Concentration values that will be used by the solver to find
    # the roots of the non-linear system.
    #
    guess = \
       {'U' : (sos, 0, 0, 0, ras),\
        'S' : simulation_guess}        # non-linear solver doesn't like
                                       # an arbitrary initial guess of lower Ras-GTP
                                       # (i.e., putting all total Ras on Ras-GDP).
                                       # Thus, we execute a numerical simulation
                                       # to estimate a better lower guess.

    for key in guess:
          
      # Computing the model steady state.
      #
      X0 = evaluator.calculate_steady_state(model.non_linear_equations, guess[key])

      if len(X0) != 0:

        # Checking correctness of the computed steady state.
        #
        is_correct = evaluator.steady_state_correctness(model.non_linear_equations, X0, 1)

        if is_correct == False:
          print('ERROR: computed steady state has negative ' +\
                'concentrations and/or violates tolerance!')
          print(X0)
          print('\n')
               

        else:
          # Storing the calculated steady state points for plotting later.
          #
          point['[SOS]'].append(sos)
          point['[GAP]'].append(gap)
          point['[RasGTP]'].append(X0[4])

          # Checking stability of steady state (array of eigenvalues).
          #
          is_stable = evaluator.check_stability(model.non_linear_equations, X0)

          is_robust = []

          if is_stable == False:
            color.append('b')
          else:
            color.append('r')

            # Checking the result robustness for stable states.
            #
            # The method returns parameters whose k-fold variation yield
            # stable steady states with high basal levels of Ras-GTP prior
            # to the bistability region (signs -/+ represent a k-fold
            # decrease/increase, respectively.
            #
            is_robust = robustness.sensitivity_analysis(model, evaluator, guess[key], X0[4])

          weak.append(is_robust)

      # else:

    # for sos in range(...

  # Storing fixed point values in tabular format
  #
  with open(dat_file, 'w') as f:

    f.write('[Total_Ras] [GEF] [SOS] [GAP] [Ras-GTP] Stable/Unstable Sensitive_parameters\n')

    for i in range(0,len(point['[SOS]'])):
      if color[i] == 'r':
        f.write("%f %f %f %f %f Stable %s\n" %\
        (ras, gef, point['[SOS]'][i], point['[GAP]'][i], point['[RasGTP]'][i], weak[i]))
      else:
        f.write("%f %f %f %f %f Unstable %s\n" %\
        (ras, gef, point['[SOS]'][i], point['[GAP]'][i], point['[RasGTP]'][i], weak[i]))

  # Plotting the final results.
  #
  plotter.plot_steady_state_graph(point, color)

if __name__ == "__main__":
  main()


# End of main.py.
#
```
