## Supplementary material for "RASGRP4 is a key factor in KRAS activation mediated by SOS in Y1 mouse tumor cell line": Source Tables and Figures: robustness.py

```
"""

robustness.py.

Module that contains classes for evaluation of result's robustness.

    Copyright (c) 2019. Marcelo S. Reis.

    This module is free software: you can redistribute it and/or modify it
    under the terms of the GNU General Public License as published by the
    Free Software Foundation, either version 3 of the License, or (at your
    option) any later version.
    ReDyMo is distributed in the hope that it will be useful, but WITHOUT
    ANY WARRANTY; without even the implied warranty of MERCHANTABILITY or
    FITNESS FOR A PARTICULAR PURPOSE. See the GNU General Public License
    for more details.
    You should have received a copy of the GNU General Public License along
    with ReDyMo. If not, see <http://www.gnu.org/licenses/>.

"""

class Robustness:

  # Constructor defines the k-fold variation (up and down) in the parameters
  # that will be applied during the sensibility analysis.
  #
  def __init__(self, k):
    self.k = k


  # This method performs an one-at-a-time sensitivity analysis, taking into
  # account a k-fold variation on all parameters.
  #
  def sensitivity_analysis(self, model, evaluator, guess, readout):

    weak_parameters = []

    for key in model.param:
   
      # k-fold increase in current model parameter.
      #    
      model.param[key] = self.k * model.param[key]      
      X_increase = evaluator.calculate_steady_state(model.non_linear_equations, guess)
      if len(X_increase) > 0 and evaluator.steady_state_correctness(model.non_linear_equations, X_increase, 1):
        is_X_increase_stable = evaluator.check_stability(model.non_linear_equations, X_increase)
  
        # 100 % tolerance over the original result
        # (it's more than enough to guarantee that no high basal
        #  levels of RasGTP is achieved during the low stable state region)
        #
        if X_increase[4] - readout > readout and is_X_increase_stable == True:
          weak_parameters.append('+' + key)
      
      # k-fold decrease in current model parameter.
      #
      model.param[key] =  (1 / (self.k * self.k)) * model.param[key]      
      X_decrease = evaluator.calculate_steady_state(model.non_linear_equations, guess)
      if len(X_decrease) > 0 and evaluator.steady_state_correctness(model.non_linear_equations, X_decrease, 1):
        is_X_decrease_stable = evaluator.check_stability(model.non_linear_equations, X_decrease)
        if X_decrease[4] - readout > readout and is_X_decrease_stable == True:
          weak_parameters.append('-' + key)
      
      # Setting current parameter value back to normal.
      #
      model.param[key] = self.k * model.param[key]

    return weak_parameters
  
  # for key in model.param:       

# End of robustness.
#
```
