## Supplementary material for "RASGRP4 is a key factor in KRAS activation mediated by SOS in Y1 mouse tumor cell line": Source Tables and Figures: simulator.py

```
"""

simulator.py.

Module that contains classes for model simulation (numeric integration).

    Copyright (c) 2019. Marcelo S. Reis.

    This module is free software: you can redistribute it and/or modify it
    under the terms of the GNU General Public License as published by the
    Free Software Foundation, either version 3 of the License, or (at your
    option) any later version.
    ReDyMo is distributed in the hope that it will be useful, but WITHOUT
    ANY WARRANTY; without even the implied warranty of MERCHANTABILITY or
    FITNESS FOR A PARTICULAR PURPOSE. See the GNU General Public License
    for more details.
    You should have received a copy of the GNU General Public License along
    with ReDyMo. If not, see <http://www.gnu.org/licenses/>.

"""

from scipy.integrate import odeint

class Simulator:

  def __init__(self):
    pass


  def simulate(self, model, y0, t):

    return odeint (model.IVP, y0, t)    


# End of simulator.py.
#
```
