## Supplementary material for "RASGRP4 is a key factor in KRAS activation mediated by SOS in Y1 mouse tumor cell line": Source Tables and Figures: steady_state.py

```
"""

steady_state.py.

Module that contains classes for evaluation of steady states.

    Copyright (c) 2019. Marcelo S. Reis.

    This module is free software: you can redistribute it and/or modify it
    under the terms of the GNU General Public License as published by the
    Free Software Foundation, either version 3 of the License, or (at your
    option) any later version.
    ReDyMo is distributed in the hope that it will be useful, but WITHOUT
    ANY WARRANTY; without even the implied warranty of MERCHANTABILITY or
    FITNESS FOR A PARTICULAR PURPOSE. See the GNU General Public License
    for more details.
    You should have received a copy of the GNU General Public License along
    with ReDyMo. If not, see <http://www.gnu.org/licenses/>.

"""

import numpy as np
import sympy as sp

import warnings

from scipy.optimize import fsolve

class SteadyState:

  # Constructor defines symbolic names of chemical species, 
  # which are used by sympy functions for stability check.
  #
  def __init__(self, symbols):
    self.X = sp.symbols (symbols, real=True)

  # Look for steady states through setting the lefthand side of ODEs
  # to zero and finding the roots of the yieled non-linear equations.
  #
  def calculate_steady_state(self, non_linear_equations, initial_guess):
    warnings.filterwarnings("error")
    try:
      solution = fsolve (non_linear_equations, initial_guess)
      return solution
    except RuntimeWarning:
      return []             # The solver could not find a solution.

  # This method verifies whether the computed state state respects
  # two properties:
  #
  # 1) No negative concentrations
  #
  # 2) The precision of the steady state is at most the number of 
  #    units specified in the 'tolerance' argument.
  #
  def steady_state_correctness(self, non_linear_equations, X, tolerance):

    Y = non_linear_equations(X)
    
    for i in range(0, len(X)):
      if X[i] < 0:                  # checking (1)
        return False
      if abs(Y[i]) > tolerance:     # checking (2)
        return False

    return True

  # Checks the steady state stability through the computation
  # of the Jacobian matrix and calculation of its eigenvalues.
  #
  def check_stability (self, non_linear_equations, X0):

    # Include the equations into a matrix, with symbolic names, 
    # in order to compute the Jacobian.
    #
    F = sp.Matrix (non_linear_equations(self.X))

    J = F.jacobian ([self.X])

    # Compute the yielded matrix J with the concentrations values.
    #
    XS = []

    for i in range (0, len(self.X)):
      XS.append((self.X[i],X0[i]))

    JS = J.subs (XS)
  
    JS_at_numpy_format = np.array (JS).astype (np.float64)

    # Compute the eigenvalues to evaluate their signal.
    #
    eigenvalues, right_eigenvectors = np.linalg.eig (JS_at_numpy_format)

    # If the steady state is stable, then for each eigenvalue its
    # real part must be negative!
    #
    for x in eigenvalues:
      if x.real >= 0:
        return False

    return True
          

# End of steady_state.py.
#
```
