## Supplementary material for "RASGRP4 is a key factor in KRAS activation mediated by SOS in Y1 mouse tumor cell line": Source Tables and Figures: test_instance_generator.py

```
"""

test_instance_generator.py.

Module to test the correctness of instance_generator.py.

    Copyright (c) 2019. Marcelo S. Reis

    This module is free software: you can redistribute it and/or modify it
    under the terms of the GNU General Public License as published by the
    Free Software Foundation, either version 3 of the License, or (at your
    option) any later version.
    ReDyMo is distributed in the hope that it will be useful, but WITHOUT
    ANY WARRANTY; without even the implied warranty of MERCHANTABILITY or
    FITNESS FOR A PARTICULAR PURPOSE. See the GNU General Public License
    for more details.
    You should have received a copy of the GNU General Public License along
    with ReDyMo. If not, see <http://www.gnu.org/licenses/>.

"""

# Path of instance_generator.py.
#
import sys
sys.path.insert(0, './src/')

import numpy as np

import pytest

from scipy.integrate import odeint

from instance_generator import *


class TestInstanceGenerator:

  def test_model_instantiation(self):

    m = RasModel()

    assert(isinstance(m, RasModel))
       

  def test_model_simulation(self):
 
    m = RasModel()

    # Simulation of 300 seconds, divided in 10001 time points.
    #
    T = np.linspace (0, 300, 10001)

    # Initial concentrations of five chemical species.
    #
    Y0 = (200, 0, 0, 0, 1000)

    X = odeint (m.IVP, Y0, T) 

    # Simulation should return, for each t in T, the corresponding
    # values for each of the five chemical species.
    #
    assert((X.size / 5) == T.size)


# End of test_instance_generator.py.
#
```
