## Supplementary material for "RASGRP4 is a key factor in KRAS activation mediated by SOS in Y1 mouse tumor cell line": Source Tables and Figures: test_steady_state.py

```
"""

test_steady_state.py.

Module to test the correctness of steady_state.py.

    Copyright (c) 2019. Marcelo S. Reis

    This module is free software: you can redistribute it and/or modify it
    under the terms of the GNU General Public License as published by the
    Free Software Foundation, either version 3 of the License, or (at your
    option) any later version.
    ReDyMo is distributed in the hope that it will be useful, but WITHOUT
    ANY WARRANTY; without even the implied warranty of MERCHANTABILITY or
    FITNESS FOR A PARTICULAR PURPOSE. See the GNU General Public License
    for more details.
    You should have received a copy of the GNU General Public License along
    with ReDyMo. If not, see <http://www.gnu.org/licenses/>.

"""

# Path of steady_state.py.
#
import sys
sys.path.insert(0, './src/')

import numpy as np

import pytest

from steady_state import *


class TestSteadyState:

  def test_stability_check(self):

    # Function to compute the steady state of the following
    # ODE system:
    #
    # dx/dt = -x^2 + 2x - xy
    #
    # dy/dt = -y + xy
    #
    def non_linear_system(arg):
      x,y = arg
      dxdt = 2 * x - x * y -x ** 2 
      dydt = x * y - y
      return (dxdt, dydt)
    
    S = SteadyState('x,y')

    # (x,y) = (0,0) is unstable
    #
    test_1 = S.check_stability(non_linear_system, (0,0))

    # (x,y) = (1,1) is stable
    #
    test_2 = S.check_stability(non_linear_system, (1,1))

    # (x,y) = (2,0) is unstable
    #
    test_3 = S.check_stability(non_linear_system, (2,0))

    assert(test_1 == False and test_2 == True and test_3 == False)
       

# End of test_steady_state.py.
#
```
