## Supplementary figures and images for "RASGRP4 is a key factor in KRAS activation mediated by SOS in Y1 mouse tumor cell line"

### FD_N1.tif

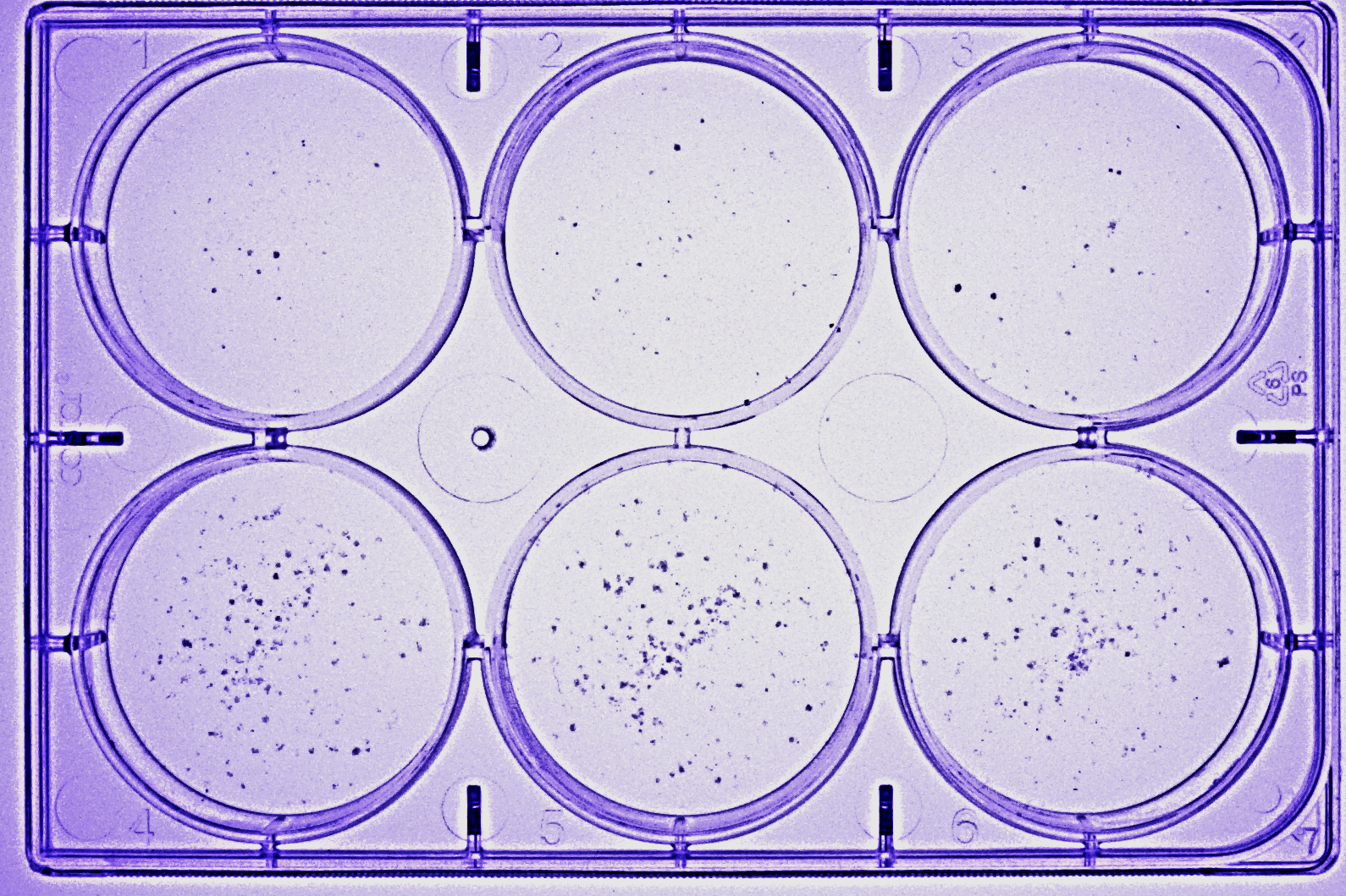

### KRAS-_N1.Tif

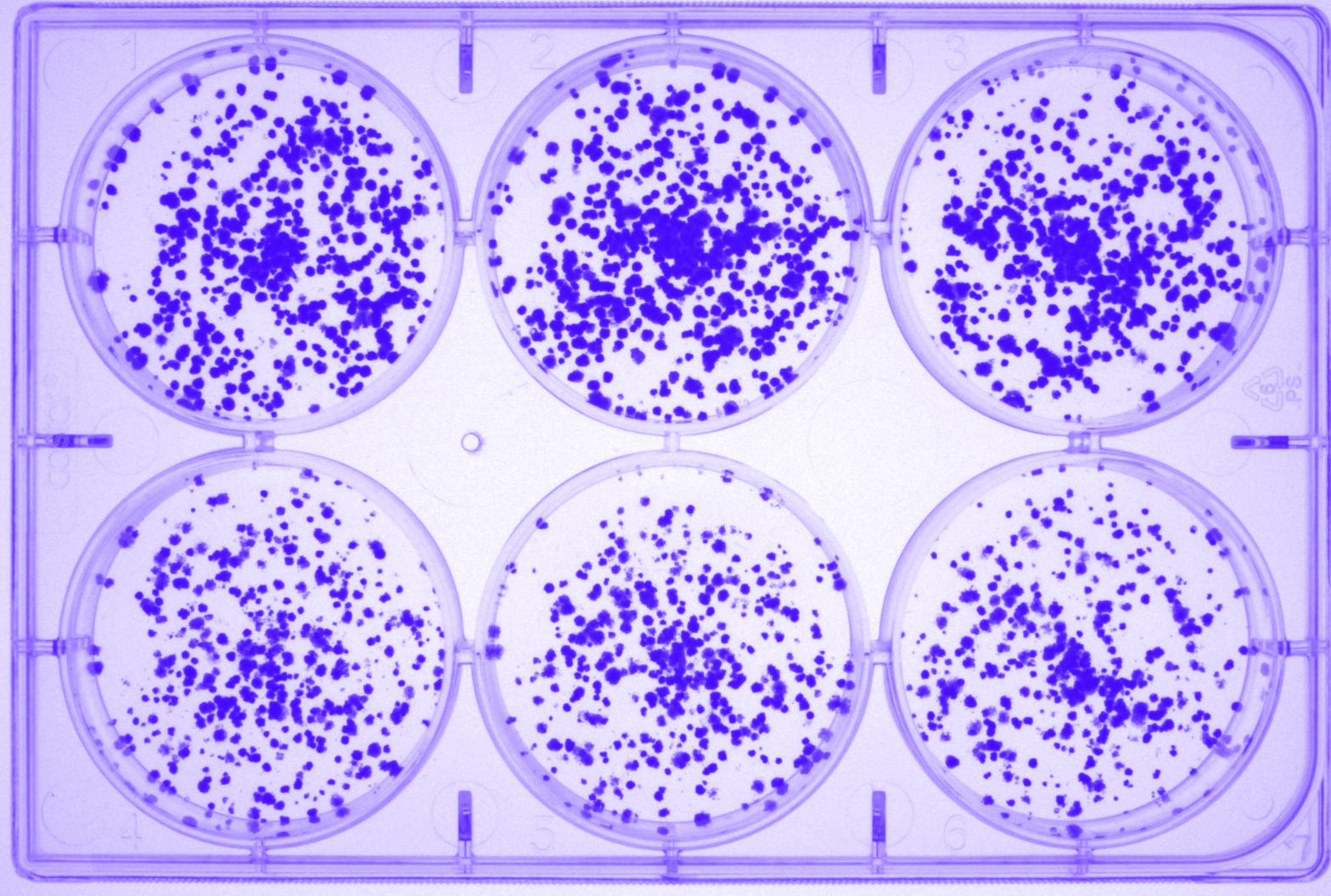

### KRAS-_N2.Tif

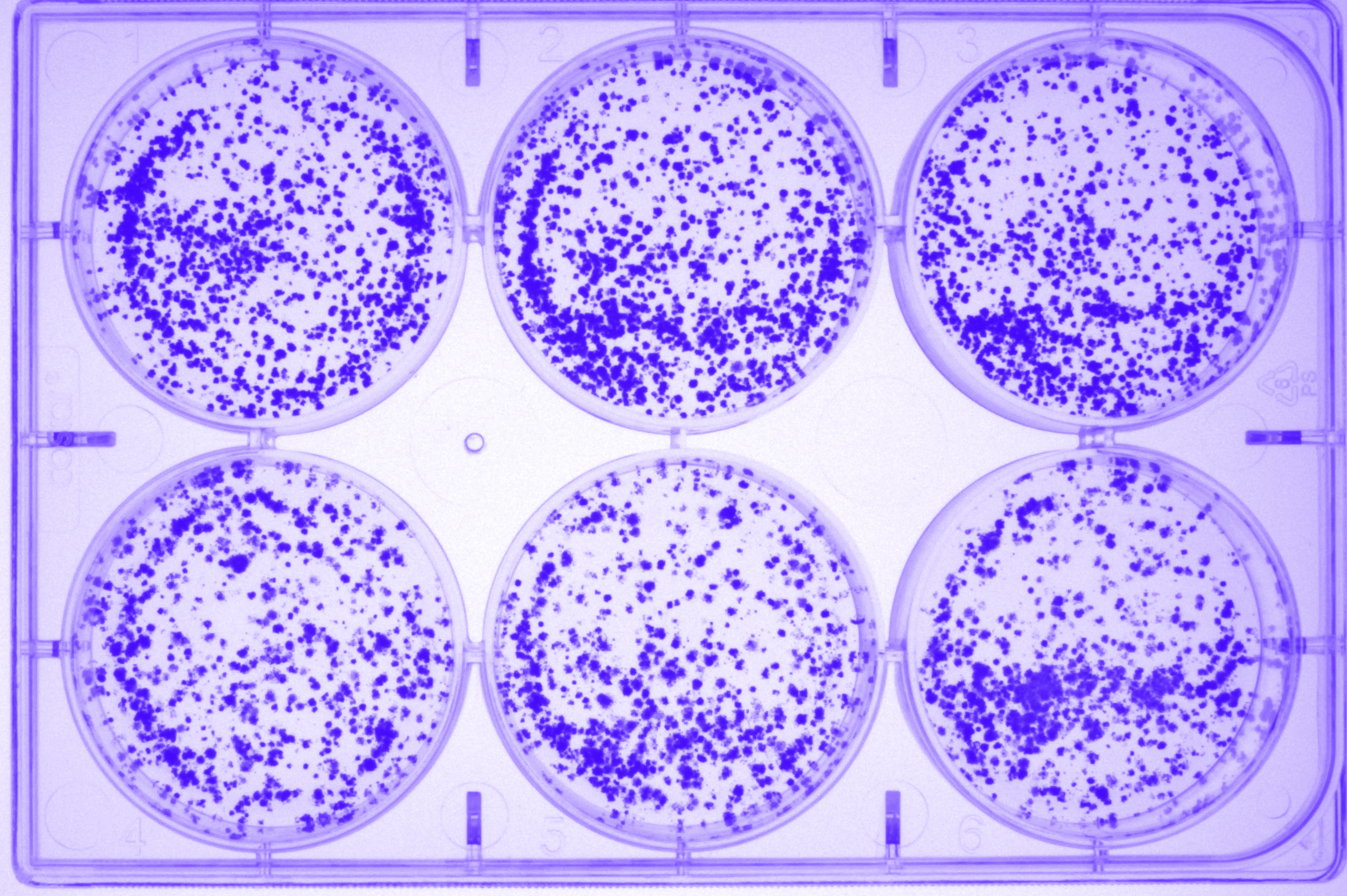

### Y1_N1.Tif

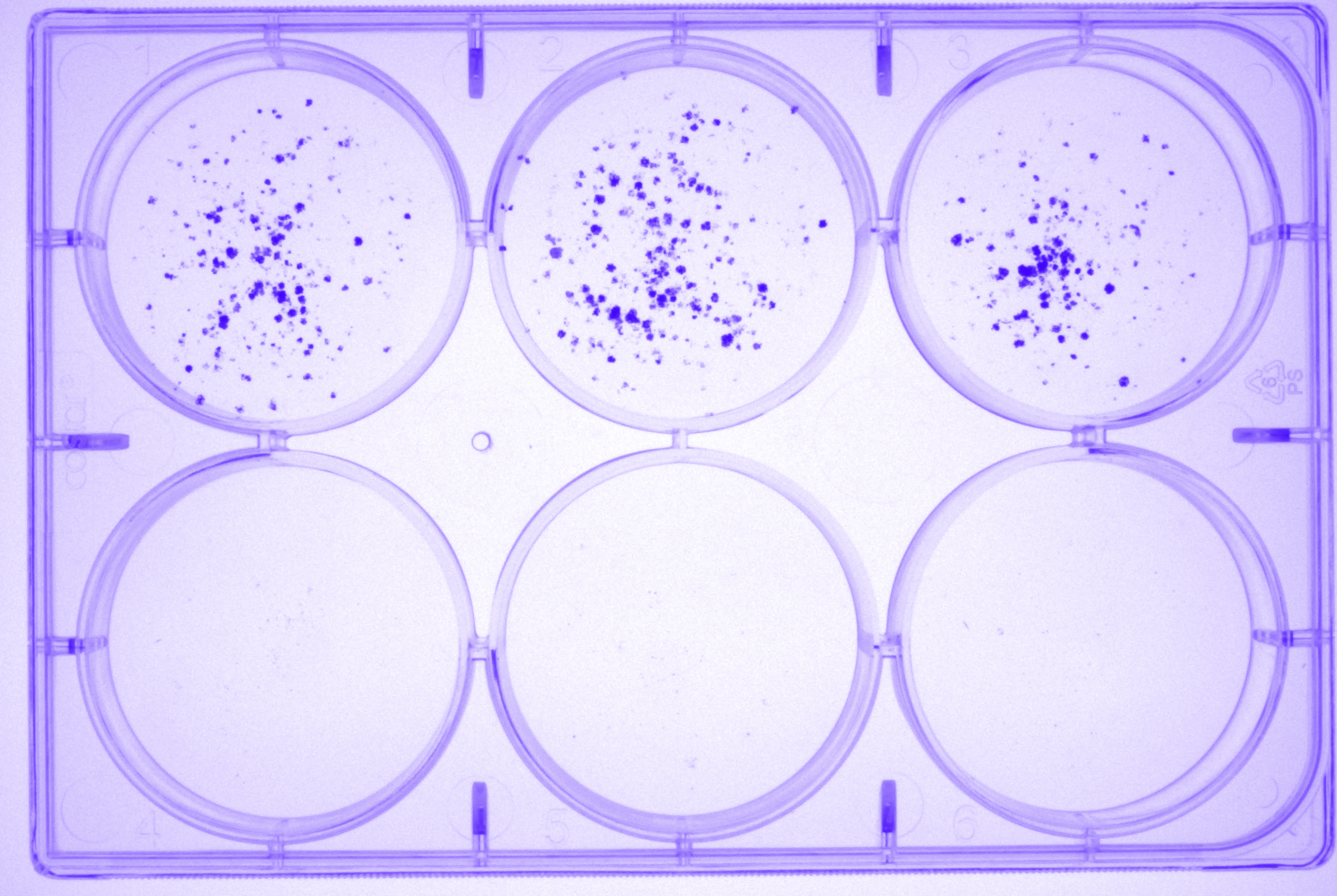

### Y1_N2.Tif

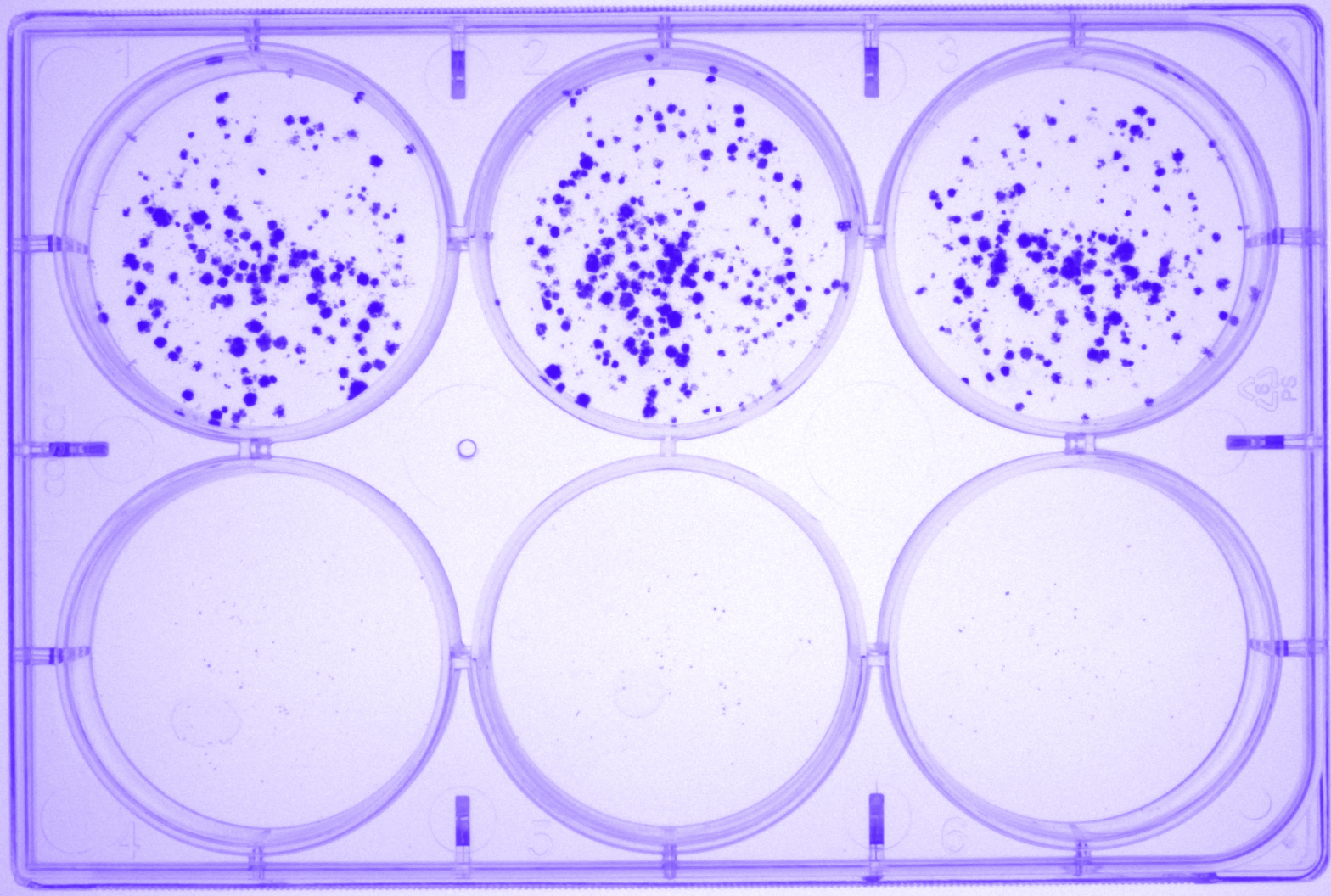

### Y1GRP4-_N1.Tif

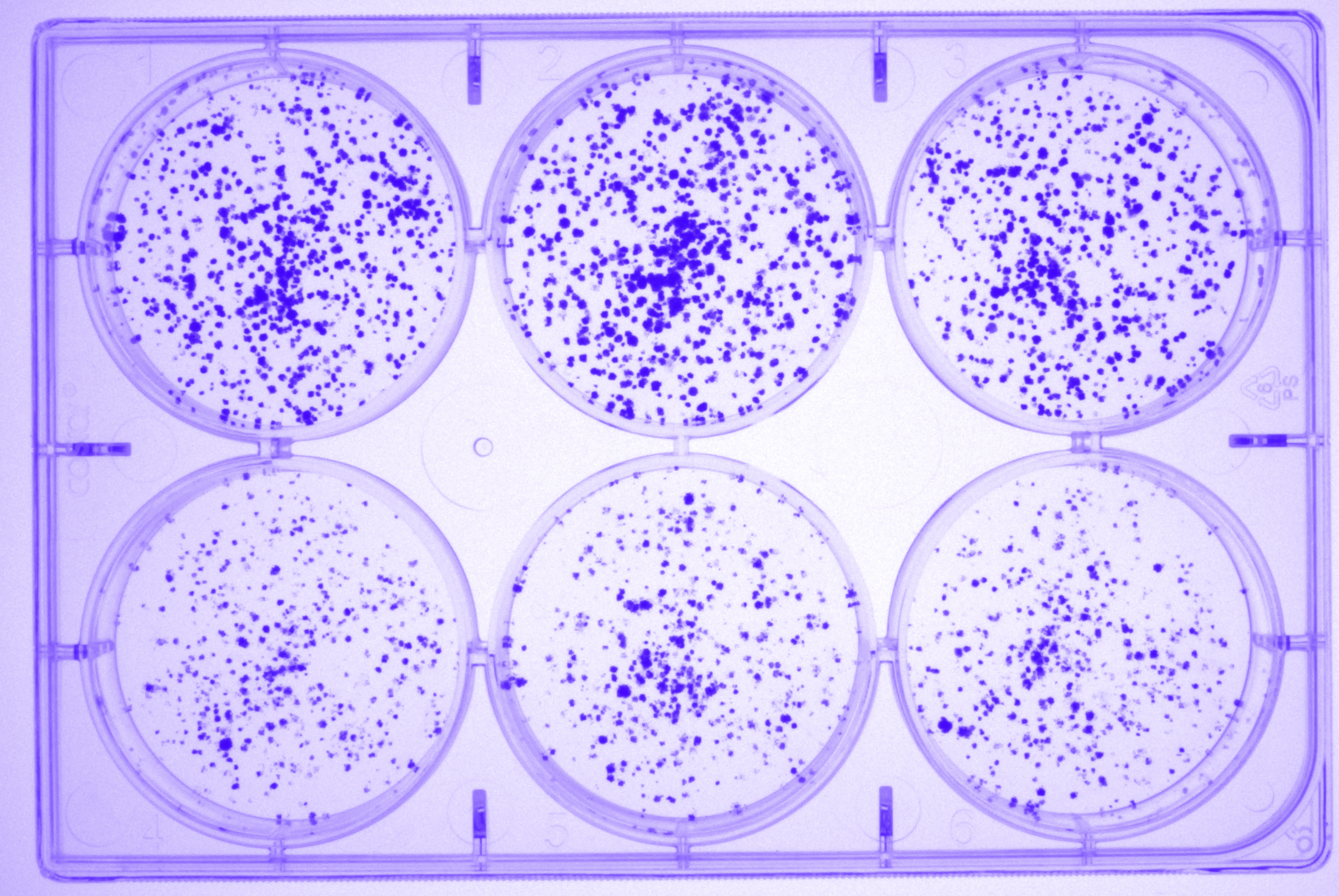

### Y1GRP4-_N2.Tif

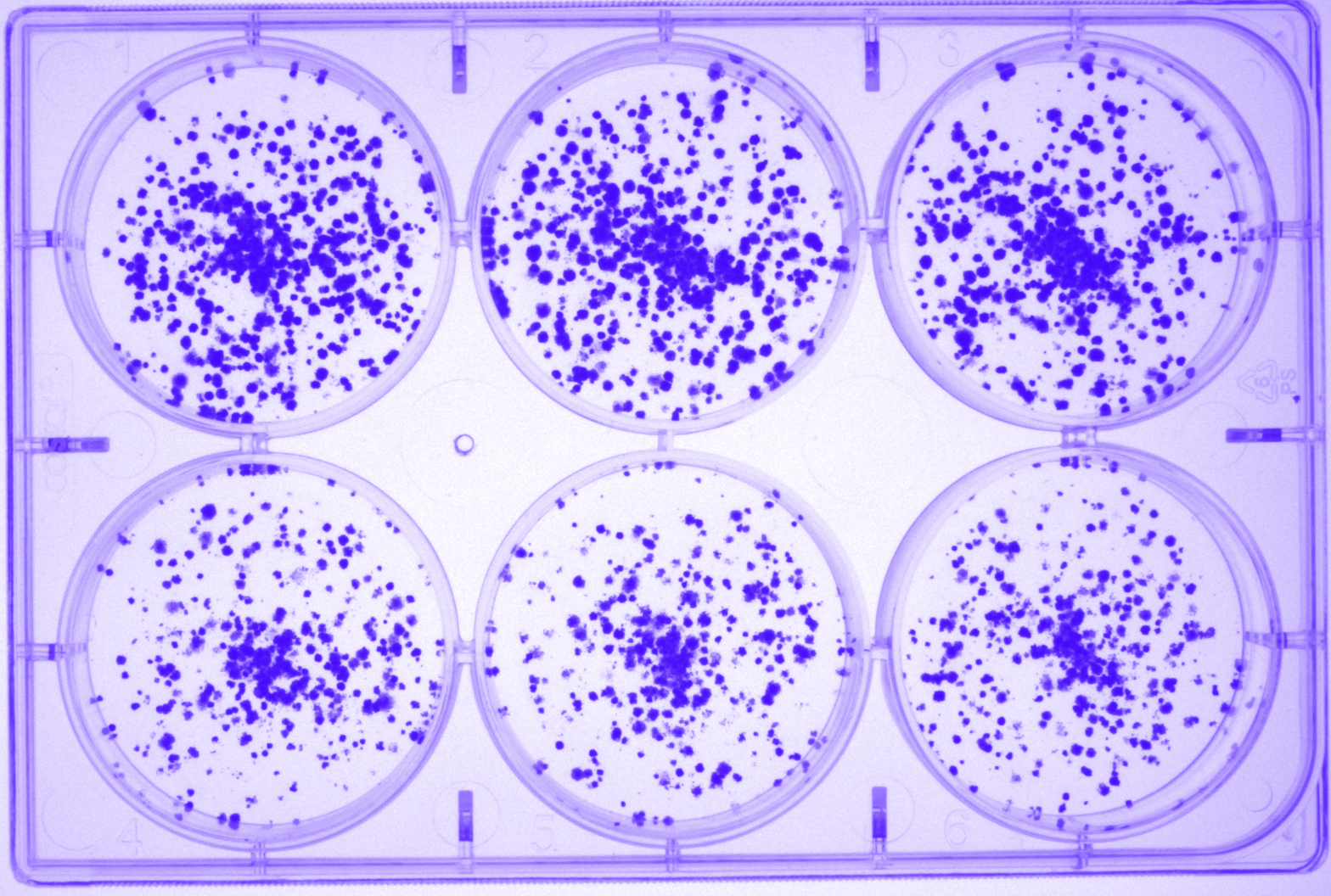
